# Conserved tyrosine Y16 acts as a phosphorylation switch controlling Ena/VASP recruitment to sites of actin assembly

**DOI:** 10.64898/2026.10.26.761804

**Authors:** Jonas Scholz, Julian Kaltenhäuser, Jerushah Emerald, Pavel Kats, Thomas Kaufmann, Anat Nitzan, Ute Curth, Klemens Rottner, Ronen Zaidel-Bar, Jan Faix

## Abstract

Ena/VASP proteins are powerful actin polymerases that accumulate at sites of active actin assembly, including lamellipodial and microspike tips, focal adhesions and pathogen surfaces, where they drive processive filament elongation. However, the mechanisms controlling their recruitment remain elusive. Here, we used cells lacking all Ena/VASP-family members to systematically test phosphomimetic and nonphosphorylatable substitutions at conserved tyrosine and serine residues implicated in subcellular targeting. Combining cell-based reconstitution with biochemical assays, we identify phosphorylation of Y16 in vertebrate Ena/VASP proteins as the decisive inhibitory modification controlling subcellular Ena/VASP recruitment, in contrast to previously proposed roles for Y39 and S157. Corresponding phosphomimetic substitutions in the *C. elegans* orthologue UNC-34 phenocopied uncoordinated locomotion of UNC-34 null worms, supporting the evolutionary conservation of this regulatory mechanism. Thus, our findings identify a conserved phosphorylation-sensitive switch at the EVH1-FP_4_ ligand interface that links EVH1-dependent targeting to dynamic control of actin assembly in adhesion, migration and pathogenesis.

## Introduction

Actin assembly at the plasma membrane provides the mechanical force required for cell protrusion, adhesion, and migration. In migrating cells, activation of Rac and the WAVE-regulatory complex (WRC) stimulates the Arp2/3 complex to generate dense, branched actin networks that drive lamellipodial protrusion ^1^. Within these networks, subsets of actin filaments can become bundled to form protruding filopodia or microspikes when they remain embedded within the lamellipodium ^2^. Sustained protrusion, however, requires continued growth of filament barbed ends at the plasma membrane. Because these ends are rapidly blocked by capping proteins (CP), their persistent elongation depends on dedicated actin polymerases such as formins and Ena/VASP proteins, which associate with growing barbed ends and protect them from capping while promoting the incorporation of additional actin monomers.

Vertebrates express three Ena/VASP-family members: VASP, Mena and Evl, which share a conserved tripartite domain architecture composed of an N-terminal Ena/VASP homology 1 domain (EVH1), a central proline-rich domain (PRD) and a C-terminal EVH2 domain ^3^. The EVH1 domain mediates the recruitment of Ena/VASP proteins to distinct subcellular sites enriched in ligands containing the canonical FPPPP (or FP_4_) motif conforming to the consensus [F/W]PXΦP (X, any amino acid; Φ, a hydrophobic residue) ^4^. Established binding partners include vinculin ^5^, zyxin ^6^, lamellipodin (Lpd, also known as Raph1) and its MRL (MIG-10/RIAM/lamellipodin) family member RIAM/PREL1 ^7,8^ as well as the *Listeria monocytogenes* surface protein ActA ^9,10^. However, EVH1-mediated interactions are not restricted to this canonical motif ^11^, as exemplified by the WRC subunit Abi, which engages the EVH1 domain through a divergent proline-rich sequence, and Testin, which employs a distinct binding interface ^12–14^. The PRD of Ena/VASP-family members binds profilin (Pfn)-actin complexes ^15^ as well as SH3-or WW-domain-containing proteins ^3^, whereas their EVH2 domains mediate tetramerization and interactions with monomeric G- and filamentous F-actin to drive actin assembly ^16–18^. Mechanistically, at steady state, the VASP tetramer remains associated with the filament barbed end through the GAB and FAB motifs of at least one EVH2 domain, while the other free EVH2 domains are able to capture actin monomers from solution and transfer them to the barbed end ^17^. Upon incorporation of a GAB-associated actin monomer, the previously barbed-end-bound EVH2 domain is thought to dissociate simultaneously from the now penultimate subunit, thereby becoming available to capture another actin monomer and initiate the next elongation cycle. Through this mode of action, the VASP tetramer is able to processively track and elongate growing barbed ends ^17,19^. Interestingly, processivity is markedly enhanced upon clustering, which enables sustained filament elongation in the presence of CP ^20,21^.

Ena/VASP proteins have been shown to localize to various sites of active actin assembly including the tips of lamellipodia, microspikes and filopodia ^22^ as well as along stress fibers (SFs) or at focal adhesions (FAs), dorsal ruffles, cell-cell contacts of epithelial cells and the surface of intracellular, pathogenic bacteria such as *Listeria monocytogenes*, as reviewed recently ^23^. More recently, VASP was shown to reside at the rim of phagocytic cups ^24,25^. Consistently, Ena/VASP proteins promote actin-driven motility across diverse systems, enhancing intracellular *Listeria* movement ^26^, bead motility in reconstituted assays ^27^, and also positively regulate migration of B16-F1 melanoma cells and fibroblasts ^28^. The latter study further revealed that Ena/VASP-deficient B16-F1 (EVM-KO) cells exhibit a perturbed lamellipodium architecture and are virtually devoid of microspikes, and are fully rescued though upon re-expression of any of the three Ena/VASP family members. Moreover, Ena/VASP-deficiency in fibroblasts also impairs the formation of FAs and leads to diminished, integrin-mediated spreading on extracellular matrix (ECM). Given these broad functions in actin-dependent adhesion, protrusion, and migration, Ena/VASP proteins have also been implicated in metastasis and carcinogenesis. Indeed, accumulating evidence links Ena/VASP proteins to a variety of cancers, including breast, gastric, pancreatic, colorectal, and lung adenocarcinomas ^29^. Notably, during the advanced phase of tumor progression, Ena/VASP proteins were found to be overexpressed, which in turn was shown to increase migration speed ^30^ and promote invasiveness ^31^. Therefore, in addition to understanding the molecular mechanism of actin assembly, it is key to elucidate the molecular mechanisms that mediate and regulate subcellular targeting of Ena/VASP-family members. However, as yet, the regulation governing subcellular Ena/VASP recruitment remains incompletely understood at the molecular level. Although EVH1-ligand interactions are known to be essential for targeting, individual EVH1-FP_4_ interactions are of relatively low affinity ^32^ and, on their own, insufficient to mediate efficient recruitment ^33^. Robust localization therefore relies on increased functional affinity through avidity, generated by multivalent interactions between tetrameric Ena/VASP proteins and clustered FP_4_ ligands. Consistent with this concept, the EVH1 domain can recapitulate effective VASP targeting only when combined with the tetramerization motif ^23^.

Moreover, it remains to be determined whether subcellular targeting is further controlled by reversible molecular switches. Unlike the WRC, which is activated by Rac to stimulate the Arp2/3 complex ^34^, or Diaphanous-related formins, which are directly regulated by Rho-family GTPases ^35,36^, Ena/VASP proteins lack canonical binding sites for Rho-family GTPases and are thus not thought to be subject to direct regulation by these GTPases. Phosphorylation therefore represents an attractive regulatory mechanism, as it is rapid and reversible and can dynamically modulate protein-protein interactions ^37,38^, making it ideally suited for the spatiotemporal coordination of protrusion, adhesion and migration. Indeed, Ena/VASP proteins are established targets of serine/threonine and tyrosine kinase signaling, and numerous phosphorylation sites have been identified by mass spectrometry and phospho-specific antibodies ^39–43^. Previous work has implicated phosphorylation of human VASP at S157 in promoting recruitment to the tip of the leading edge and FAs ^44^, whereas phosphorylation of Y39 in the VASP-EVH1 domain by c-Abl (also known as Abl1) was reported to interfere with FA targeting ^41^. Intriguingly, the EVH1 domain contains a second surface-exposed tyrosine residue, Y16, which, like Y39, is conserved across VASP, Mena, and Evl and has been found to be phosphorylated in multiple cell types ^42^, including HEK-293T ^45^, HT1080 ^46^, non-small cell lung cancer cell lines ^47^, Jurkat T cells ^48^, and B-lymphocytes ^49^. However, its relevance has not yet been analyzed.

Of note, previous conclusions regarding the effects of phosphorylation were derived from studies performed in wild-type (WT) cells or cells only partially depleted of endogenous Ena/VASP proteins. Given, however, that Ena/VASP family members can hetero-oligomerize ^50^, residual endogenous paralogs may influence the behavior of ectopically expressed mutants and thereby complicate interpretation of results. Therefore, here we used genuine Ena/VASP-deficient B16-F1 mouse melanoma cells and fibroblasts as reconstitution systems to define how conserved phosphorylation sites regulate Ena/VASP targeting. Importantly, we also validated our results *in vivo*, using the model organism *C. elegans*. Our findings support a model in which phosphorylation at Y16 disrupts multivalent EVH1-dependent interactions with clustered FP_4_ ligands, thereby shutting off recruitment of Ena/VASP proteins to sites of active actin assembly, providing a universal switch for the spatiotemporal regulation of these powerful actin polymerases.

## Results

### Phosphorylation of S153 in murine VASP, corresponding to S157 in human VASP, does not affect subcellular targeting

Phosphorylation of S157 in human VASP has been reported to drive the subcellular targeting of the protein to the leading edge or FAs (Fig. 1a) ^39,44,51^, while phosphorylation of Y39 has been assumed to prevent recruitment of VASP to these sites ^41^. To reassess the proposed role of S157 phosphorylation in the subcellular targeting of VASP in genuine Ena/VASP-deficient B16-F1 mouse melanoma cells (EVM-KO), we first generated the phosphomimetic mouse VASP (mVASP) mutant S153D, in which S153, corresponding to S157 in human VASP, was replaced with aspartic acid to mimic constitutive phosphorylation. Substitution of the same residue with alanine (S153A) rendered this site nonphosphorylatable. We then expressed these mVASP variants fused to EGFP in EVM-KO cells and assessed the subcellular localization of these constructs and of endogenous vinculin as a marker for FAs by immunolabeling. However, we found no evidence for an improved localization of the phosphomimetic mutant S153D to subcellular sites, neither to the leading edge nor to FAs, since the nonphosphorylatable VASP mutant S153A localized virtually identically to the S153D mutant at the tip of the leading edge (Lp) and in FAs (Fig. 1b). Thus, in contrast to previous assumptions, phosphorylation at this site evidently has no noticeable effect on the subcellular targeting of VASP.

**Fig. 1:**
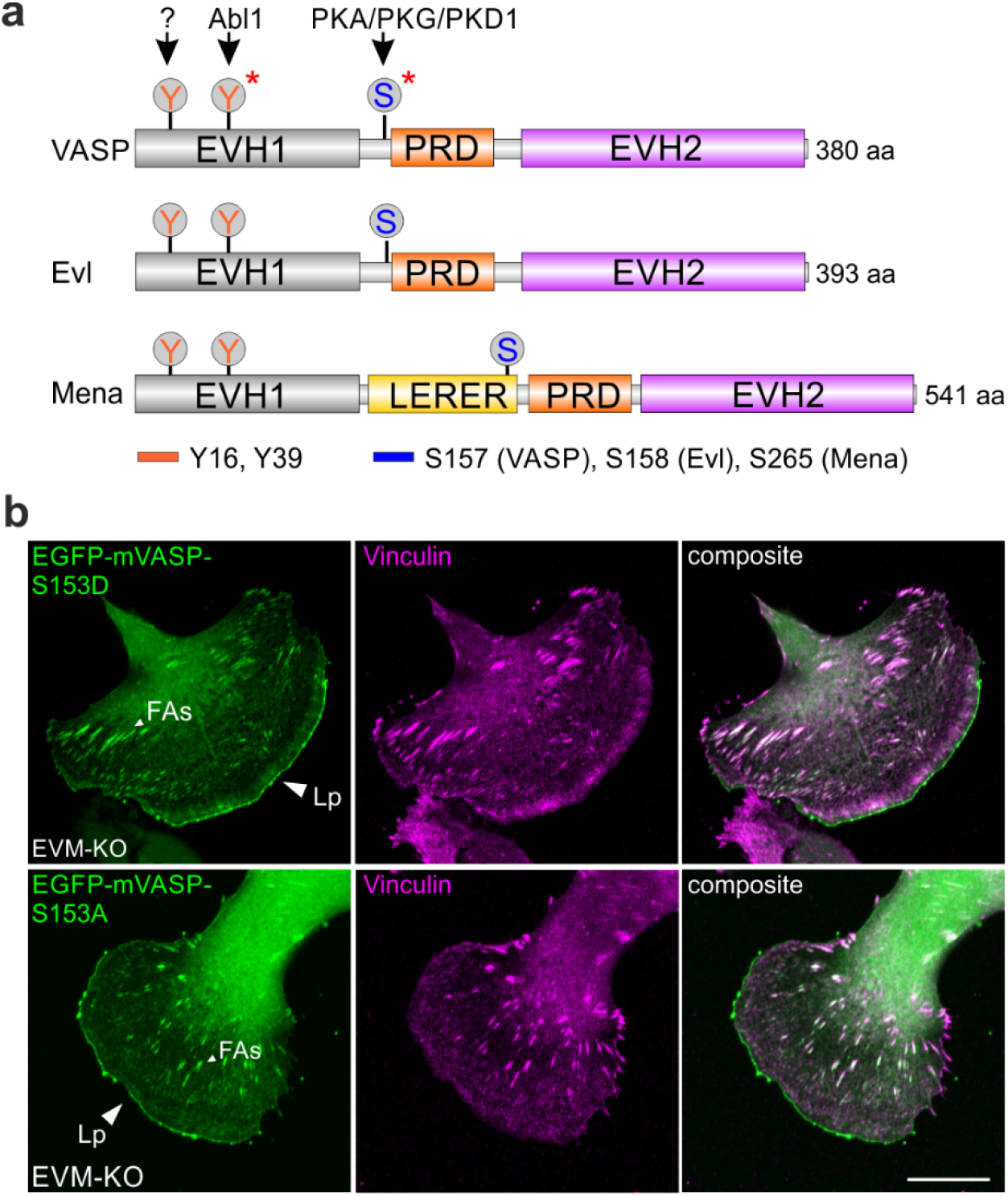
Phosphorylation of S153 in murine VASP does not affect subcellular targeting of the protein to the lamellipodial leading edge and to FAs. **a** Scheme of Ena/VASP proteins with highlighted tyrosine (Y) and serine (S) phosphorylation sites linked to subcellular targeting. The numbering corresponds to the major human protein isoforms, with the kinases indicated above by black arrows. Residues Y16 and Y39 are fully conserved across all three Ena/VASP paralogs compared to the corresponding serine residues. Red asterisks indicate phosphorylated Y39 and S157 of human VASP, which have been reported to regulate subcellular targeting ^41,44^. Abbreviations: EVH1, Ena/VASP homology domain 1; PRD, proline-rich domain; LERER, low complexity region harboring LERER repeats; EVH2, Ena/VASP homology 2. **b** Phosphorylation of S153 in murine VASP (mVASP), corresponding to S157 in human VASP, does not affect subcellular targeting of the protein to the tips of lamellipodia (Lp) and embedded clusters representing microspike tips or to focal adhesions (FAs). Representative images of Ena/VASP-deficient EVM-KO cells transiently expressing EGFP-tagged phosphomimetic mVASP-S153E or nonphosphorylatable mVASP-S153A while migrating on laminin. Cells were fixed and stained with GFP nanobodies and antibodies against the FA marker vinculin. Note the colocalization with VASP in FAs, but the lack of vinculin at the lamellipodium tip, as previously established ^101^. Scale bar, 20 μm. Abbreviations: Lp, lamellipodium tip; FAs, focal adhesions.

### Phosphorylation of Y16, but not of Y39, blocks accumulation of VASP at the tips of lamellipodia

The major tyrosine phosphorylation site implicated in subcellular targeting of human VASP is Y39 within the EVH1 domain (Fig. 1a). In wild-type NIH 3T3 fibroblasts, Y39 was reported to be critical for zyxin binding and FA targeting ^41^. However, high-throughput mass spectrometry data indicate that Evl is predominantly phosphorylated at Y16 rather than Y39 ^42^, although this apparent difference may also reflect differential detectability of the respective phosphopeptides ^52^. Both residues are conserved across all three Ena/VASP family members in mammals, suggesting potential roles of both residues in subcellular targeting. Notably, however, the crystal structure of the Mena-EVH1 domain bound to an ActA-FP_4_ peptide revealed that only Y16 resides directly at the FP_4_-binding interface, whereas Y39 is located in a distant β-strand (Fig. 2a) ^53^, suggesting that the latter may potentially regulate FP_4_-ligand binding indirectly through an allosteric mechanism or directly through an additional binding site, as previously reported for zyxin ^54^. To experimentally reassess the functional relevance of Y16 and Y39 phosphorylation for subcellular VASP localization, each residue was substituted with either phenylalanine or glutamate to generate nonphosphorylatable and phosphomimetic mutants, respectively. Stable cell lines were subsequently generated using EVM-KO cells, followed by FACS-mediated cell sorting to obtain populations with comparable expression levels of the respective EGFP-mVASP variants, both among each other and relative to endogenous VASP levels in B16-F1 control cells (Fig. 2b). We then asked whether microspikes could be reformed upon expression of phosphomimetic (Y16E and Y39E) and the nonphosphorylatable (Y16F and Y39F) VASP mutant proteins fused to EGFP in EVM-KO cells. To our surprise, but in full agreement with the distant position of Y39 to the FP_4_-ActA peptide in the crystal structure, expression of phosphomimetic mutant Y39E fully rescued microspike formation with VASP accumulating in clusters at their tips and the periphery of the leading edge, demonstrating that this VASP mutant was functional (Fig. 2c-e; Movie 1). Accordingly, its localization was indistinguishable from the reconstitution experiment with the wild-type (WT) protein or the nonphosphorylatable Y39F variant. In sharp contrast, stable expression of the phosphomimetic Y16E mutant at near-physiological levels largely prevented VASP accumulation in clusters and at the front of the leading edge, thereby impairing microspike formation, whereas the nonphosphorylatable Y16F variant showed appropriate VASP localization and efficiently rescued microspike formation. Consistent with the major targeting defect, mVASP-Y16E was typically distributed throughout the entire lamellipodial actin network rather than displaying the crisp leading-edge accumulation characteristic of VASP-WT and Y16F, with little or no enrichment at the leading-edge tip. This distribution resembled that of previously described VASP variants comprising only the actin-binding EVH2 domain in Mena/VASP-deficient MV^D7^ cells ^55^. Similar localization defects were also observed for the corresponding Y16E mutants of the other two Ena/VASP paralogues, Mena and Evl (Supplementary Fig. 1). Consistent with the localization data, only EVM-KO cells reconstituted with mVASP-Y16E failed to rescue the pronounced 2D migration defect of the triple-knockout cells on laminin ^28^, whereas all other VASP variants restored migration rates to levels observed in B16-F1 control cells (Fig. 2f). Then, we calculated the mean square displacement (MSD) of the cells to assess their effective directional movement. As expected, EVM-KO cells reconstituted with mVASP-Y16E displayed MSD values similar to those of the parental EVM-KO cells and markedly lower than those of all other reconstituted cell lines plus B16-F1 control (Supplementary Fig. 2a). Finally, we asked whether locking VASP in a non-phosphorylated state would increase protrusion persistence. To this end, we used time-lapse phase-contrast imaging to determine the duration of lamellipodial protrusions. Interestingly, however, we observed only a slight, statistically insignificant increase in lamellipodial persistence (Supplementary Fig. 2b), suggesting that loss of negative VASP regulation alone is insufficient to prevent termination of lamellipodial protrusions.

**Fig. 2:**
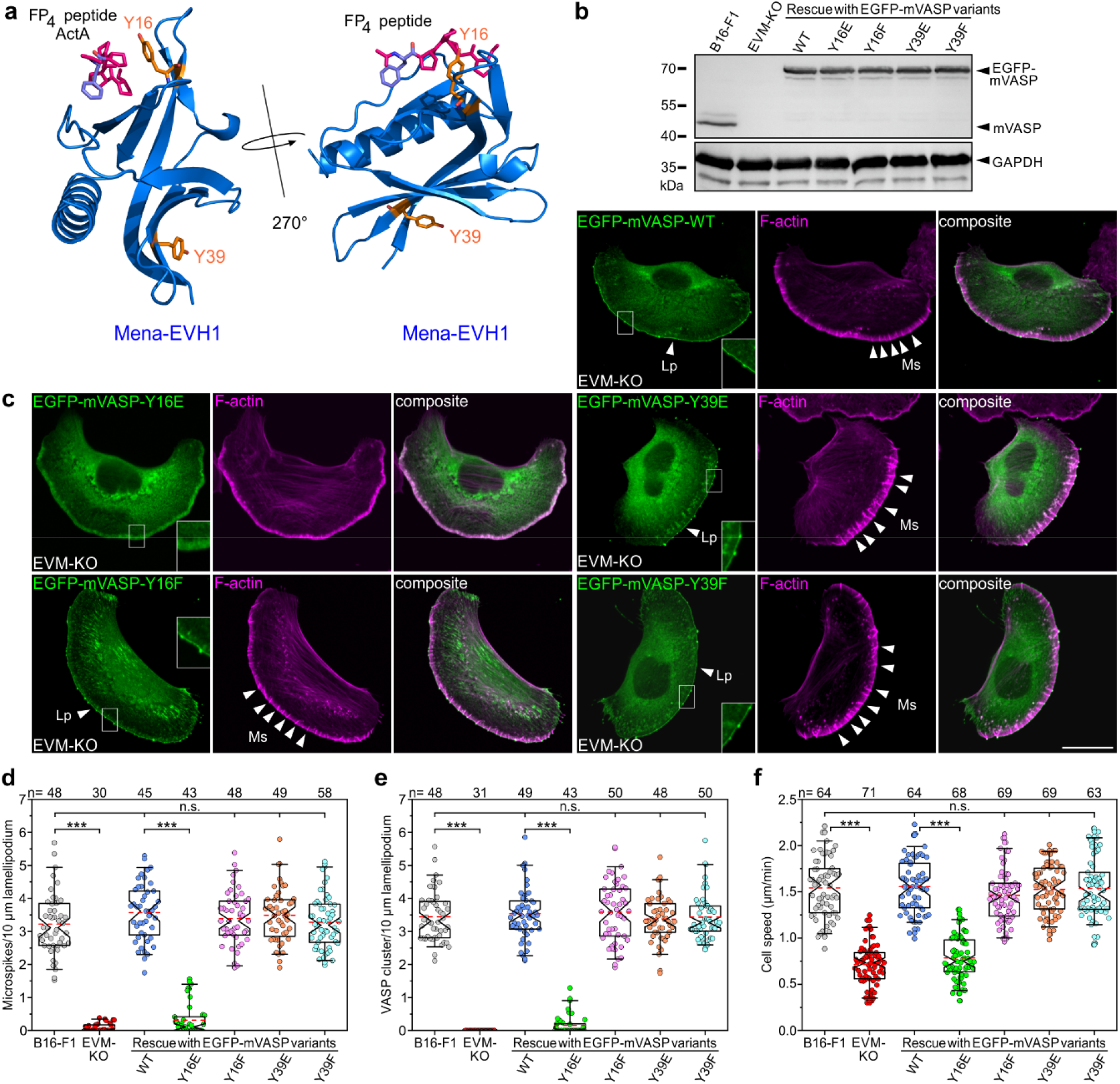
Phosphorylation of Y16, but not of Y39, blocks accumulation of VASP at the tips of lamellipodia and microspikes. **a** Ribbon diagram of the crystal structure of murine Mena-EVH1 domain in complex with the FP_4_ peptide from *L. monocytogenes* ActA (modified from RCSB PDB: 1evh.pdb) ^53^. Note that in the complex, surface exposed Y16 directly faces the FP_4_ motif, whereas Y39 is located further away on the opposite side of the structure. **b** Stable expression of EGFP-tagged mVASP-variants as indicated in EVM-KO cells. GAPDH blot is shown as loading control. **c** Phosphomimetic Y16E, but not of Y39E, blocks accumulation of VASP at the tips of lamellipodia and microspikes. Representative images of EVM-KO cells stably expressing EGFP-tagged phosphomimetic or nonphosphorylatable mVASP-variants as indicated while migrating on laminin. Cells were fixed and stained with GFP nanobodies and phalloidin for the actin cytoskeleton. Brightest point projections of confocal stacks are shown. Scale bar, 20 μm. Insets, enlarged images of boxed regions. Note that microspike formation is not rescued in cells expressing EGFP-mVASP-Y16E. Abbreviations: Lp, lamellipodium tip; Ms, microspikes. **d** Quantification of microspike density in lamellipodia. **e** Quantification of VASP clusters. **f** Quantification of cell speed on laminin. (**d-f**) Boxes in box plots indicate 50% (25 to 75%) and whiskers 90% (5 to 95%) of all measurements, with dashed red lines depicting the means. Medians are highlighted by indentation of boxes. Kruskal-Wallis test with Dunn’s multiple comparison test. *n*, number of cells. Results are pooled data from at least three biologically independent replicates. \*\*\**P* < 0.001. n.s., not significant.

### Phosphorylation of Y16 blocks accumulation of VASP at focal adhesions

VASP is a well-established component of the actin-regulatory layer of FAs, where it is recruited predominantly in a mechanosensitive manner by zyxin and other FP_4_ motif-containing proteins ^56,57^. In general, fibroblasts constitute a better model system for the analysis of canonical FAs than B16-F1 melanoma cells migrating on laminin. To test directly whether VASP recruitment to FAs is also regulated by phosphorylation at Y16 and/or Y39, we ectopically expressed our phosphomimetic and nonphosphorylatable mVASP constructs in MVE-KO fibroblasts lacking all three Ena/VASP family proteins ^28^ (Fig. 3a). Of note, colocalization analysis of mVASP with the FA marker vinculin revealed that, among all variants tested, only the phosphomimetic Y16E mutant failed to localize to FAs (Fig. 3b). To determine whether this effect extends to other Ena/VASP family members, we next examined the FA localization of EGFP-tagged mEvl and mMena variants. Similar to mVASP, all mEvl variants were strongly enriched at vinculin-positive FAs, except for Y16E, which again showed no detectable FA localization (Supplementary Fig. 3a). In contrast, mMena-Y16E retained partial FA localization, potentially due to its unique LERER repeats previously shown to interact directly with integrin α5 ^58^, whereas all other mMena variants were strongly enriched at FAs (Supplementary Fig. 3b). Together, these findings establish Y16 as a critical determinant of Ena/VASP recruitment to FAs, with Mena-specific LERER repeats providing an alternative targeting mechanism.

**Fig 3:**
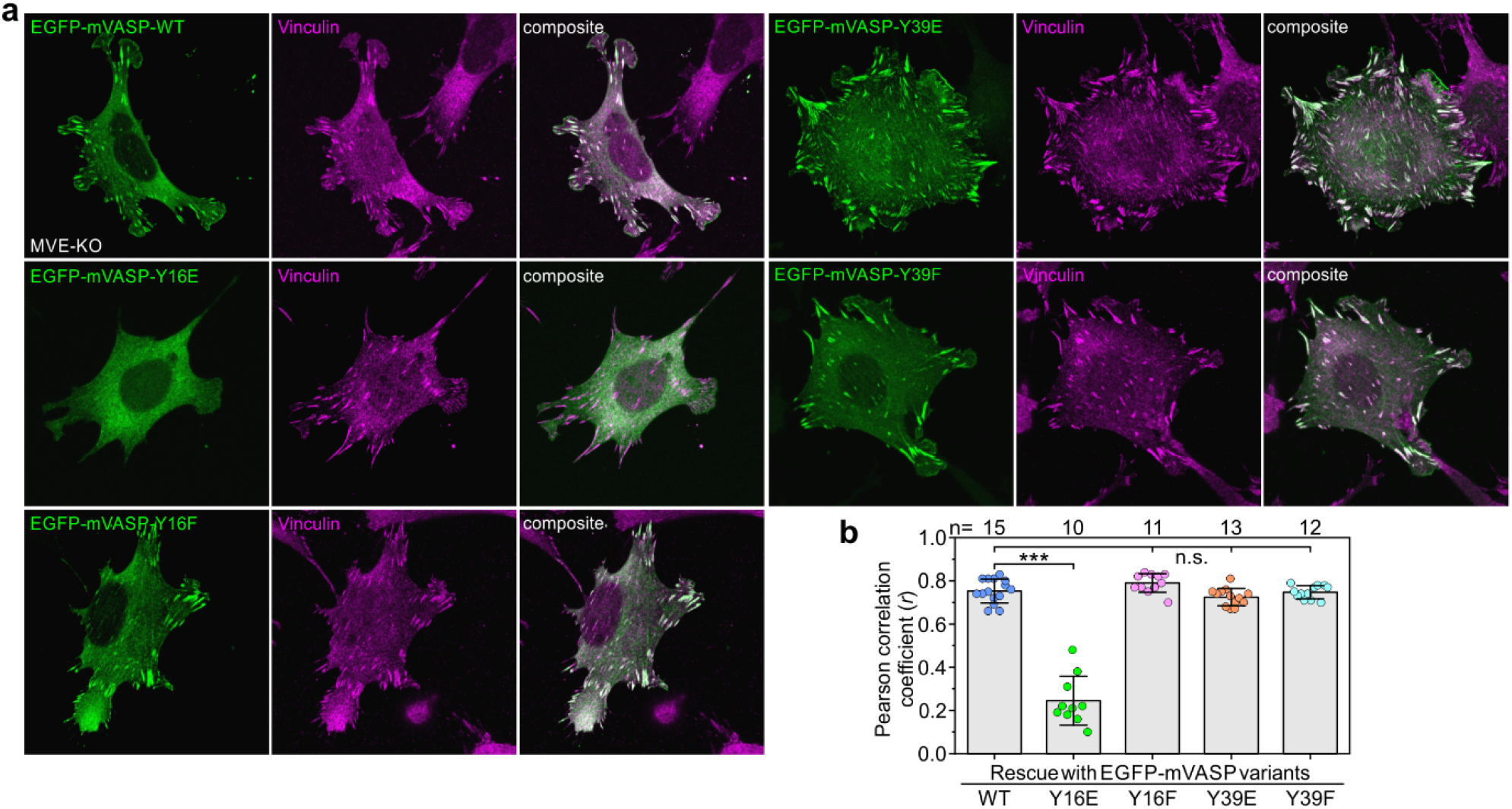
Phosphorylation of Y16, but not of Y39, blocks accumulation of VASP at focal adhesions. **a** Representative images of Ena/VASP-deficient MVE-KO fibroblasts ^28^ transiently expressing EGFP-tagged mVASP variants as indicated. With the exception of the phosphomimetic Y16E mutant, all other mVASP variants tested co-localized with vinculin in FAs. Brightest point projections of confocal stacks are shown. Scale bar, 20 μm. **b** Assessment of VASP and vinculin colocalization. Respective intensity distributions in regions excluding the leading edge were used to calculate the corresponding Pearson correlation coefficients. Bars represent arithmetic means ± SD. One-way ANOVA with Tukey’s multiple comparison test. *n*, number of cells. Results are pooled data from at least three biologically independent replicates. \*\*\**P* < 0.001. n.s., not significant.

### VASP Y16E, but not Y39E, disrupts ActA binding and actin assembly on Lpd-derivatized beads

Next, we asked whether the localization defects of the Y16 variants are directly caused by perturbed FP_4_-ligand binding. To address this question, we characterized Ena/VASP-FP_4_ interactions using complementary *in silico* modeling based on existing structural data and *in vitro* assays using the established FP_4_ ligands ActA from *L. monocytogenes* and Lpd. First, we performed a series of umbrella sampling (US) molecular dynamics (MD) simulations to calculate and compare the dissociation free energies Δ*G*_diss_ for the unbinding of an ActA-derived FP_4_ peptide from the WT Mena-EVH1 domain, the phosphomimetic Y16E variant, and the EVH1 domain phosphorylated at Y16 (pY16). To this end, we used the 1.8 Å crystal structure of the Mena-EVH1 domain bound to the consensus proline-rich ActA peptide ‘FPPPPT’ (PDB: 1EVH) ^53^, in which Y16 directly faces the bound peptide (Fig. 4a). Notably, Δ*G*_diss_ was markedly reduced for both EVH1 variants compared with WT, corresponding to decrease in binding affinities of approximately two and six orders of magnitude, respectively, and thus indicating substantially weakened interactions with the FP_4_ peptide, particularly for EVH1-pY16 (Fig. 4b and c). Thus, the clearly impaired binding of the phosphomimetic Y16E variant supports the use of Ena/VASP-Y16E variants as valid phosphomimetic models in cellular assays and is consistent with the mislocalization of these constructs observed in our functional reconstitution experiments in cells. Interestingly, individual EVH1-ligand interactions are relatively weak, with affinities typically lying in the micromolar range ^32^. Thus, VASP tetramerization is thought to promote multivalent binding, thereby increasing avidity (effective affinity) and residence time at subcellular sites to promote efficient subcellular targeting ^23^. We therefore sought to validate our computational findings in the context of full-length, tetrameric murine VASP (mVASP_4_). To this end, we employed sedimentation velocity analytical ultracentrifugation (AUC) to assess the interaction of mVASP-WT, mVASP-Y16E, and mVASP-Y39E with an ActA fragment (aa 248-389) encompassing all four FP_4_ motifs ^32^. We found that addition of ActA to mVASP_4_-WT at a substoichiometric 0.5:1 molar ratio shifted approximately half of the tetramer population from 3.3 S to 4.3 S, while the remaining fraction stayed unbound (Fig. 4d). Increasing ActA concentrations up to an eightfold molar excess over mVASP_4_ did not further increase the sedimentation coefficient, consistent with formation of a high-affinity complex between ActA and mVASP_4_ (Fig. 4d). Titration experiments further confirmed a 1:1 stoichiometry between mVASP_4_ and ActA (Supplementary Fig. 4a). Importantly, analysis of the VASP variants revealed that even at an eight-fold molar excess of ActA, no binding was detected for phosphomimetic mVASP_4_-Y16E, whereas mVASP_4_-Y39E binding to ActA was indistinguishable from VASP_4_-WT (Fig. 4e and f; Supplementary Fig. 4b and c).

**Fig 4:**
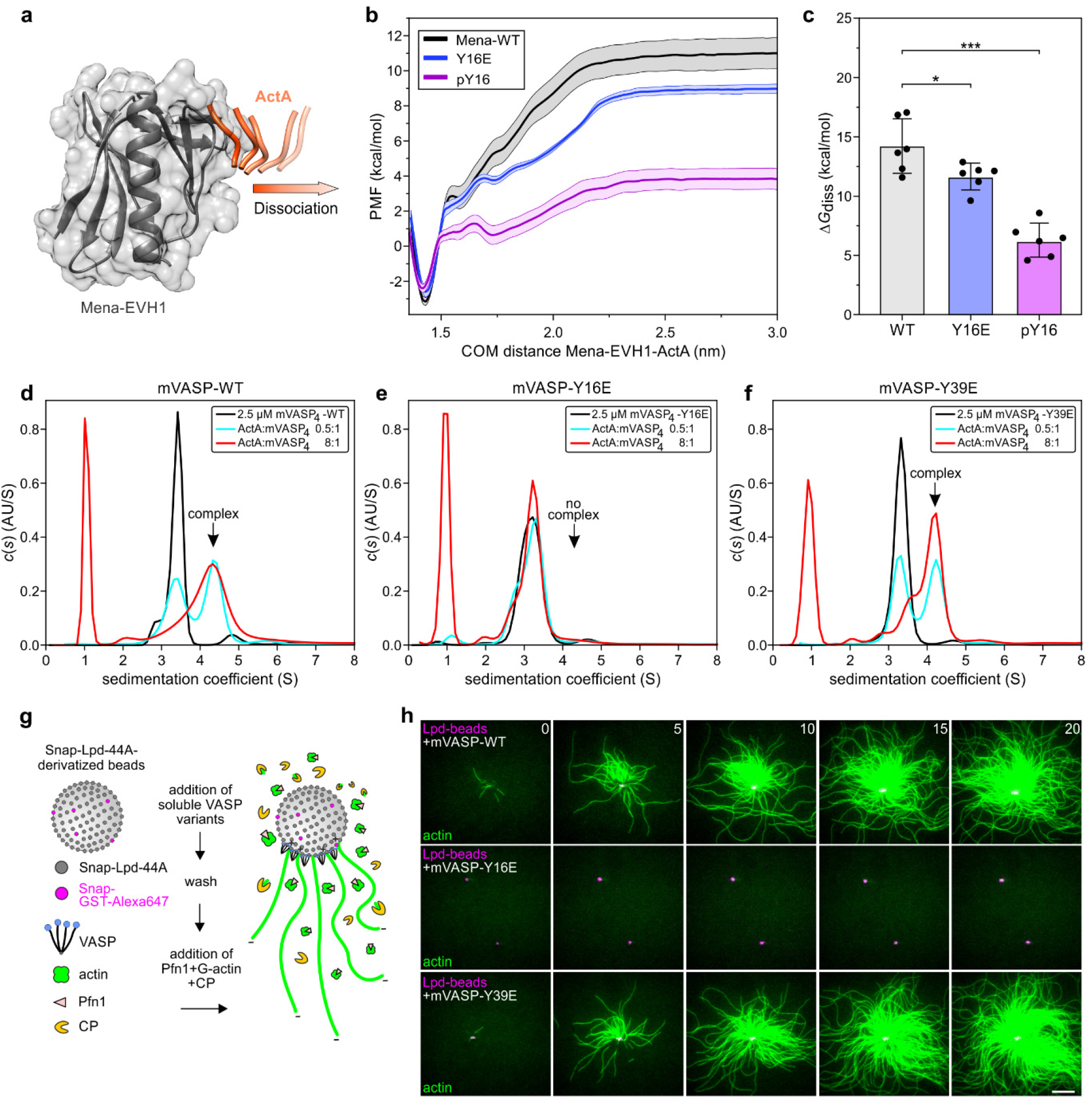
Phosphorylation at VASP Y16 disrupts ligand binding and downstream actin assembly. **a** Schematic representation of the pulling MD simulations. The crystal structure of the Mena-EVH1 domain (grey) bound to a short ActA FP_4_ peptide (PDB: 1EVH) ^53^, served as the starting model for the umbrella-sampling simulations (see Materials and Methods for details). The direction of the pulling force applied to the ActA peptide is indicated by the arrow. **b** Potential of mean force (PMF) curves for ActA dissociation from WT, phosphomimetic Y16E, and phosphorylated Y16 (pY16) Mena-EVH1 domains. Each curve represents the average from six independent calculations. Curves are shown as mean ± SEM. COM, center-of-mass. **c** Dissociation free energies Δ*G*_diss_, quantified as the difference between the maxima and minima of the PMF curves shown in (**b**). Data are shown as mean ± SD. **d** Sedimentation velocity analytical ultracentrifugation analysis of the interaction of ActA (aa 248-389) and the mVASP tetramer (mVASP_4_). Note shift to higher *s*-values of the VASP tetramer in presence of ActA confirming their interaction. Of note, even an 8-fold molar excess of ActA (red line) did not noticeably increase the *s*-value of the complex as compared to the 0.5:1 mixture (cyan), strongly suggesting that the VASP tetramer forms a high-affinity 1:1 complex with ActA. Results for the experiments performed with 2.5 µM of the VASP tetramer and 1.25 or 20 µM ActA, respectively, are shown. **e-f** Under comparable experimental conditions, no complex was detected with the phosphomimetic Y16E mutant, whereas Y39E formed a complex indistinguishable from VASP-WT. **g** Scheme of TIRF-microscopy assays analyzing VASP-mediated actin assembly on SNAP-Lpd^850–1250^(44A)-derivatized beads in the presence of CP and Pfn1. **h** Representative still images from time-lapse TIRF movies using 3 μM actin (10% ATTO488-labeled) together with 25 nM CP and 5 µM Pfn1 on Lpd^850–1250^(44A)-derivatized beads pre-incubated with the soluble mVASP variants as indicated. Time is indicated in min. Scale bar, 10 µm. The SNAP-capture beads were visualized by the addition of trace amounts (5%) of ATTO647-labelled GST-SNAP. Note processive actin-filament elongation with mVASP-WT and phosphomimetic mutant Y39E, and lack of actin assembly with phosphomimetic mutant Y16E.

To expand our analysis and assess the relevance of VASP tyrosine phosphorylation at Y16 *versus* Y39 for its recruitment to and stimulation of processive actin assembly at surfaces *in vitro*, we employed SNAP-capture magnetic beads derivatized with the SNAP-tagged dimeric Lpd fragment GCN4-Lpd^850–1250^(44A), which harbors all six FP_4_ motifs but lacks additional actin-binding sites ^13,59^. We then monitored actin assembly in the presence of soluble mVASP-WT and both phosphomimetic mutants, CP, and profilin 1 (Pfn1) in real time at the single-filament level using multicolor total internal reflection fluorescence (TIRF) microscopy (Fig. 4g). Consistent with previous experiments, preincubation of the beads with mVASP-WT ^13^ or phosphomimetic mVASP-Y39E resulted in robust actin assembly on their surfaces in the presence of CP, consistent with functional VASP clustering, whereas no detectable actin assembly was detected on beads incubated with mVASP-Y16E (Fig. 4h, Movie 2). Taken together, these data strongly suggest that phosphorylation at Y16 selectively disrupts EVH1-FP_4_ interactions, thereby preventing VASP recruitment and VASP-driven actin assembly at ligand-coated surfaces, whereas phosphorylation at Y39 has no detectable effect.

### Phosphomimetic VASP-Y16 disrupts Listeria actin comet-tail formation and motility

Bacterial pathogens frequently hijack the host actin cytoskeleton at multiple stages of infection, exploiting actin remodeling to promote cellular entry, intracellular motility, and dissemination to neighboring cells. A prominent example is *L. monocytogenes*, which exploits the host actin machinery for intracellular motility and cell-to-cell spread through its surface protein ActA, a bacterial actin nucleation-promoting factor that directly activates the Arp2/3 complex while simultaneously recruiting Ena/VASP proteins via its clustered FP_4_ motifs to promote efficient actin assembly and comet-tail formation ^60–62^. Consistent with previous work, VASP accumulated prominently and asymmetrically at the surface of *L. monocytogenes*, coinciding with the formation of a long actin comet tail, whereas the Ena/VASP-deficient EVM-KO cells analyzed here supported the formation of short actin stubs only (Fig. 5a; Movie 3), closely resembling phenotypes previously observed in MV^D7^ fibroblasts lacking Mena and VASP expression ^63^. Importantly, reconstitution of EVM-KO cells with EGFP-mVASP-WT rescued comet tail formation (Fig. 5b). To assess the function of VASP phosphorylation on Y16/39, we then expressed the entire collection of phosphomimetic and nonphosphorylatable mutants. Of note, only reconstitution with mVASP-Y16E failed to restore the formation of canonical actin comet tails, whereas all remaining variants, including mVASP-Y39E, fully rescued this process, resulting in comet-tail lengths comparable to those observed in B16-F1 WT cells (Fig. 5c and d; Movie 4). In line with this observation, mVASP-Y16E also failed to restore *Listeria* motility, in contrast to mVASP-WT (Supplementary Fig. 5). Nevertheless, we still observed minute but clearly detectable accumulation of mVASP-Y16E at the bacterial surface. To test whether this residual recruitment might be mediated, at least in part, by association with actin, we expressed a combinatorial mutant lacking G- and F-actin binding domains (ΔGAB and ΔFAB, respectively) on top of Y16E. Interestingly, mVASP-Y16EΔGAB/ΔFAB was virtually abolished in recruitment to the bacterial pole, whereas the corresponding actin-binding-deficient mVASP-WT variant (ΔGAB/ΔFAB) remained prominently enriched at this site, albeit unable to rescue comet tails (Fig. 5e and f). Thus, although interactions with actin can apparently contribute to the residual accumulation at bacterial surfaces seen with mVASP-Y16E, this mechanism appears inefficient to restore comet-tail formation. In addition, they also confirm those direct actin interactions to be obligatory for Ena/VASP function at the bacterial pole, even if recruited in an EVH1-FP_4_-mediated fashion. Together, these data are fully consistent with FP_4_-mediated VASP recruitment being obligatory for driving processive actin assembly downstream of Arp2/3 complex activation in this system.

**Fig 5:**
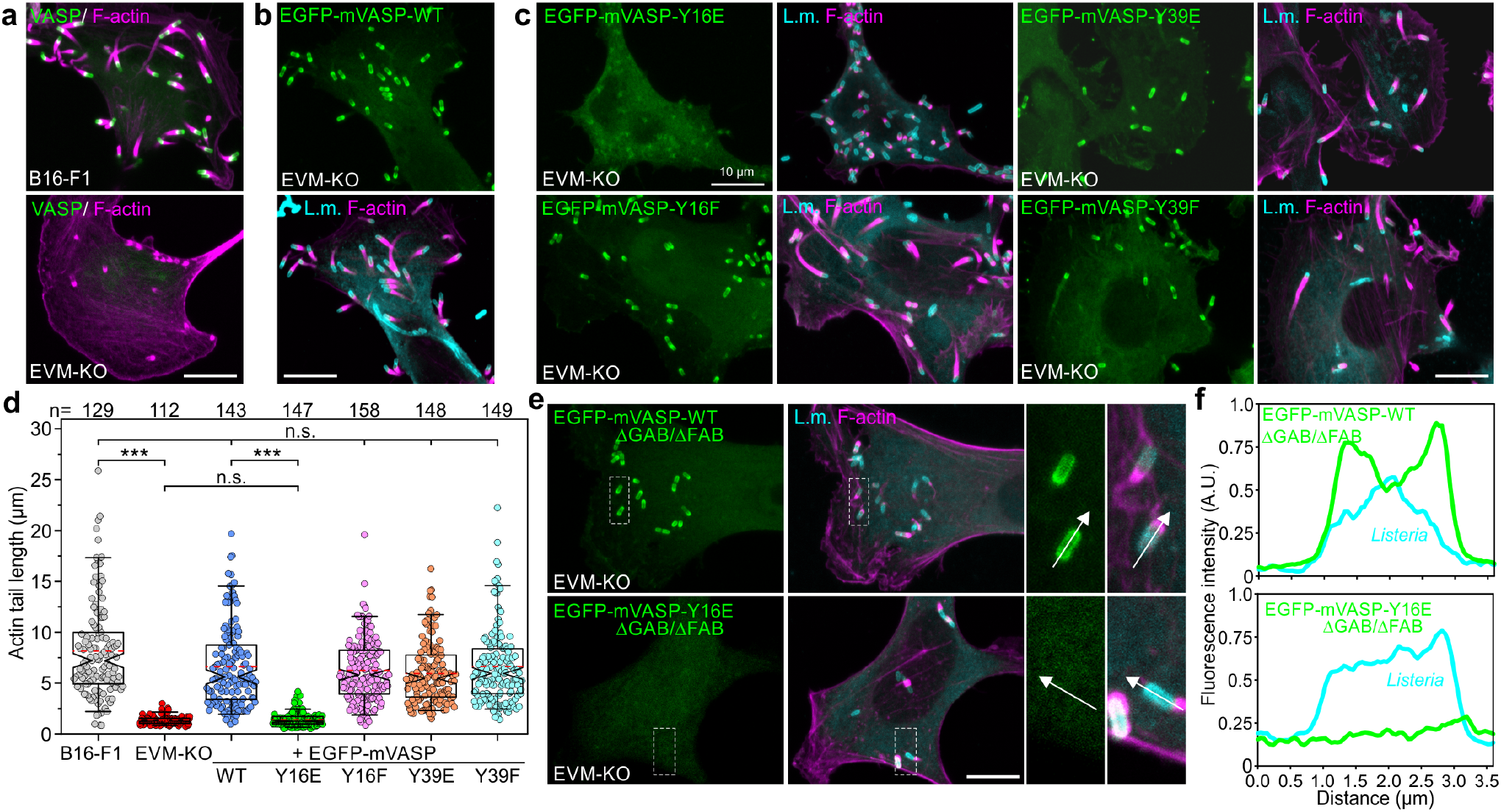
Phosphorylation of VASP at Y16 perturbs *Listeria*-associated localization and actin comet tail assembly. **a** Representative images of *Listeria* actin comet tails in infected B16-F1 WT and EVM-KO cells stained for endogenous VASP (green) and the actin cytoskeleton (magenta). Note the short, stubby comet tails in the absence of Ena/VASP proteins. **b** Reconstitution of EVM-KO cells with EGFP-tagged mVASP-WT fully restores *Listeria* comet tail formation. *Listeria* are shown in cyan. **c** Representative images of EVM-KO cell lines stably expressing the EGFP-tagged mVASP variants as indicated. With the exception of the Y16E mutant, all other mVASP variants markedly accumulated at the surface of *L. monocytogenes* and promoted the formation of long actin comet tails. Nevertheless, residual amounts of EGFP-mVASP-Y16E still accumulated at the bacterial surface. Brightest point projections of confocal stacks are shown. **d** Quantification of actin comet tail length from cells as shown in (**c**). Boxes in box plots indicate 50% (25 to 75%) and whiskers 90% (5 to 95%) of all measurements, with dashed red lines depicting the means. Medians are highlighted by indentation of boxes. Kruskal-Wallis test with Dunn’s multiple comparison test. *n*, number of actin comet tails. Results are pooled data from at least three biologically independent replicates. \*\*\**P* < 0.001. n.s., not significant. **e** Representative images of EVM-KO cells reconstituted with EGFP-tagged mVASP-WT or mVASP-Y16E variants lacking the G- and F-actin-binding sites. Loss of these modules, which are essential for filament-barbed end tracking ^17^, virtually abolished recruitment of mVASP-Y16E to the bacterial surface, in contrast to mVASP-WT. Enlarged images of boxed regions are shown on the right. **f** Intensity profiles of VASP and the *Listeria* surface along white arrows as depicted in (**e**). (**a-c, e**) Scale bars, 10 μm.

### Phosphomimetic UNC-34-Y17E in C. elegans results in uncoordinated locomotion, phenocopying UNC-34 loss of function

Interestingly, the EVH1 domain of the single *C. elegans* Ena/VASP ortholog UNC-34 ^64,65^ contains only one tyrosine residue, Y17, corresponding to Y16 in mammalian Ena/VASP proteins, while lacking an equivalent corresponding to Y39 (Fig. 6a). We therefore used the nematode *C. elegans* to test whether regulatory tyrosine phosphorylation within the EVH1 domain is conserved across evolutionarily distant species. To this end, we employed CRISPR/Cas9-mediated genome editing to replace Y17 of endogenous UNC-34 with either the phosphomimetic Y17E or the nonphosphorylatable Y17F variant. We then compared the locomotion of WT and mutant L4 larvae on agar plates seeded with bacterial lawns using time-lapse imaging (Fig. 6b; Movie 5). Subsequent analysis revealed that the phosphomimetic UNC-34-Y17E mutant exhibited a severe uncoordinated locomotion phenotype closely resembling that of UNC-34-null mutants ^66,67^, whose impaired movement has been attributed to defective neuronal migration ^64^. This phenotype was evident from sequential overlays of whole-animal contours over time and analyses of worm curvature (Fig. 6c and Supplementary Fig. 6). In contrast, UNC-34-Y17F animals displayed coordinated locomotion closely resembling that of WT controls, albeit at a reduced speed (Fig. 6d and e). Additionally, we found that synchronized L4 larvae lacking UNC-34 or expressing UNC-34-Y17E were considerably smaller than larvae expressing nonphosphorylatable UNC-34-Y17F or UNC-34-WT (Fig. 6b). Given their pronounced locomotion defects, this reduced body size may result from impaired access to the bacterial food source and consequently reduced nutrient uptake, leading to slower larval growth. Thus, consistent with the data obtained in mammalian cells, these findings identify Y17 as a critical regulatory site whose phosphomimetic modification severely compromises UNC-34-dependent locomotion in *C. elegans*.

**Fig. 6:**
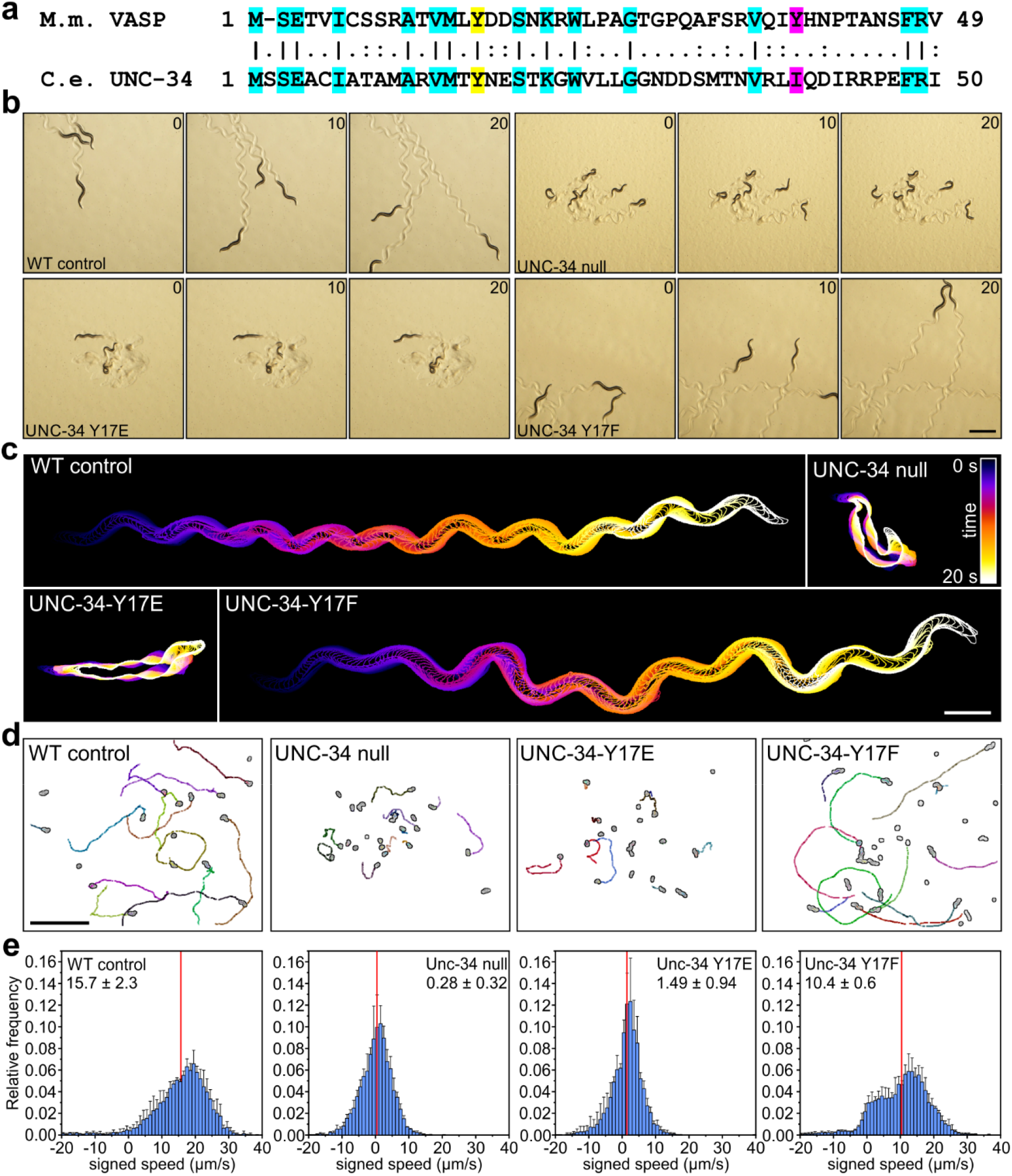
The critical function of the first tyrosine residue is conserved in *C. elegans*. **a** Alignment of the N-terminal regions of the EVH1 domains of murine VASP and *C. elegans* UNC-34. Identical residues are highlighted in cyan, except for the conserved tyrosine residue corresponding to Y16 in murine VASP and Y17 in UNC-34, which is shown in yellow. Notably, Y39 of murine VASP (magenta), which is conserved across all mammalian Ena/VASP proteins, is absent from UNC-34. **b** Still images from brightfield time-lapse movies of WT, UNC-34-null, UNC-34-Y17E, and UNC-34-Y17F L4 larvae migrating on bacterial lawns. Time is indicated in seconds. Scale bar, 500 µm. These data correspond to Movie 5. **c** Time-color-coded outlines of the animals illustrate the dramatic difference in their motile behavior. Scale bar, 500 µm. **d** Tracks of migrating L4 larvae on bacterial lawns. Scale bar, 5 mm. **e** Quantification of worm speed from experiments as shown in (**d**). Signed speed indicates forward (positive values) or backward (negative values) movement, with values near zero reflecting pauses or minimal movement. Histograms show the relative frequency of speeds (bin width: 1 µm/s), presented as mean ± SD across three independent biological replicates (*n* = 3), with at least seven worms analyzed per replicate. Red lines indicate the mean of the biological replicate means, with corresponding values reported as mean ± SD.

## Discussion

In this study, we identified a conserved tyrosine residue within the EVH1 domain of Ena/VASP-family members as critical determinant of their localization and function. By combining genetic reconstitution of genuine Ena/VASP-deficient cells to directly assess phenotypic rescue with biochemical analyses and molecular dynamics simulations, here we demonstrate that phosphomimetic substitution of VASP-Y16, but not Y39, severely compromises interactions with FP_4_-containing ligands. Consequently, VASP-Y16E fails to accumulate efficiently at lamellipodial tips, FAs and intracellular *Listeria*, resulting in a failure to rescue microspike formation, cell migration and actin comet-tail assembly. Moreover, introduction of the corresponding Y17E substitution into the endogenous *unc-34* locus causes severe locomotion defects in *C. elegans*, highlighting the evolutionary conservation of this critical residue. Together, our findings support a model in which phosphorylation of Y16 regulates Ena/VASP-dependent actin assembly by controlling spatial recruitment rather than directly inhibiting actin polymerase activity.

Interestingly, previous work implicated phosphorylation of human VASP-S157 in promoting recruitment to lamellipodia and FAs ^44^. However, phosphomimetic and nonphosphorylatable substitutions of the corresponding murine residue S153 had no noticeable effect on VASP localization in our Ena/VASP-deficient experimental system. While the reasons for the discrepancy with previous reports remain unclear, they may reflect the presence of residual endogenous Ena/VASP proteins in the earlier study. Nevertheless, we note that our findings are not unexpected, given that S153 lies outside both the EVH1 domain and the tetramerization motif, which are critical for proper subcellular targeting ^23^.

Conversely, the strikingly different consequences of Y16E and Y39E substitutions can be explained by the distinct structural environments of these residues. Previous studies established that Y16, together with W23 and F79, forms an aromatic triad of surface-exposed side chains that converge in the folded, globular EVH1 domain to create a hydrophobic recognition surface for proline-rich ligands ^4,40,53^. Introducing a negatively charged residue at this position would therefore be expected to interfere with ligand recognition. Indeed, our molecular dynamics simulations and AUC experiments revealed pronounced destabilization of the EVH1-ActA interaction upon either phosphomimetic substitution or phosphorylation of Y16, providing a molecular explanation for the defects observed experimentally. By contrast, Y39 is positioned on the opposite face of the EVH1 domain and does not directly contribute to the canonical FP_4_-binding groove. Instead, previous structural and biochemical analyses identified a secondary EVH1 ligand-binding surface involving Y39 that contributes to the recognition of multivalent zyxin ligands ^54^. Phosphomimetic substitution of Y39 was reported to disrupt this secondary interaction while largely preserving canonical ligand recognition, thereby altering the binding mode and stoichiometry of EVH1-zyxin complexes. In our AUC experiments using ActA, however, Y39E formed ActA-VASP complexes with the same 1:1 stoichiometry as VASP-WT.

Our biochemical findings further highlight the importance of multivalent EVH1-ligand interactions for stable Ena/VASP recruitment. Whereas tetrameric VASP-WT and VASP-Y39E readily formed complexes with an FP_4_-containing ActA fragment, VASP-Y16E failed to interact even at elevated ligand concentrations. Interestingly, ActA binding caused a pronounced increase in the sedimentation coefficient of VASP, which is unlikely to be explained by the relatively small increase of about 10% in molecular mass alone. Rather, this observation suggests a substantial change in hydrodynamic properties, consistent with compaction of the otherwise highly flexible VASP tetramer ^17,20^. Given that a single ActA molecule can simultaneously engage up to four EVH1 domains ^32^, such compaction could arise from multivalent engagement of the four EVH1 domains within a single VASP tetramer by closely spaced FP_4_ motifs. This interpretation is in line with the relatively weak affinities of individual EVH1-FP_4_ interactions and the proposed role of multivalency in promoting stable Ena/VASP recruitment to sites of actin assembly ^23,68^. The functional importance of this interaction was further demonstrated in our bead-based *in vitro* assays, in which VASP-WT and VASP-Y39E efficiently promoted actin assembly at surfaces functionalized with an FP_4_-containing lamellipodin fragment, whereas VASP-Y16E failed to be recruited to the bead surface from solution to initiate actin polymerization. Thus, defective ligand recruitment provides a compelling explanation for the inability of Y16E to support localized actin assembly, without requiring a direct impairment of its intrinsic polymerase activity.

The cellular consequences of this targeting defect were particularly apparent at the leading edge of migrating cells. Recent work showed that lamellipodin and Abi contribute to leading-edge Ena/VASP accumulation and turnover, whereas unconventional myosin X is essential for VASP clustering and microspike formation ^13,69^. In this context, the diffuse distribution of VASP-Y16E throughout the lamellipodial actin network, together with its failure to accumulate in microspike-associated clusters, emphasizes the requirement for an intact EVH1 recognition interface. Although residual association with the actin network may occur through the EVH2 domain, this interaction alone appears insufficient to establish the spatial organization necessary for proper lamellipodium protrusion and microspike assembly. Moreover, the inability of Y16E to restore efficient cell migration suggests that appropriate positioning of Ena/VASP proteins is essential for coordinating protrusive actin dynamics with cell movement. With the exception of Mena, whose integrin-binding LERER repeats are sufficient for FA targeting ^58^, similarly pronounced targeting defects to these sites were observed for the Y16E variants of VASP and Evl, demonstrating that the relevance of this tyrosine residue extends beyond the lamellipodial leading edge.

The failure of VASP-Y16E to support *L. monocytogenes* motility further reinforces the essential role of EVH1-mediated recruitment in actin-based of protrusion and pushing forces, which relies in this case on the interaction of host Ena/VASP proteins with the FP_4_-containing bacterial surface protein ActA. Notably, the faint residual accumulation of Y16E on *Listeria* was virtually abolished upon deletion of G- and F-actin-binding motifs within EVH2, suggesting that limited recruitment may occur more indirectly through association with actin. However, this residual recruitment was unable to support comet-tail formation. These findings therefore distinguish a mere actin-dependent, downstream association from productive, EVH1-mediated upstream targeting and emphasize that recruitment through FP_4_-containing ligands is critical for efficient, Ena/VASP-dependent actin assembly coinciding or complementing obligatory Arp2/3 complex activation in this system ^60^.

The functional importance of this conserved tyrosine extends to *C. elegans*, where phosphomimetic substitution of the equivalent residue Y17 in UNC-34, the sole nematode Ena/VASP ortholog lacking the Y39 counterpart, caused severe locomotion defects resembling those of UNC-34-null animals, supporting an evolutionarily conserved role of Y16 in Ena/VASP function. Of note, animals carrying the nonphosphorylatable Y17F substitution remained largely coordinated, although their locomotion speed was reduced. This modest defect may reflect the inability of Y17F to undergo reversible phosphorylation, potentially limiting the dynamic regulation of UNC-34 activity required for optimal locomotion. Although the cellular defects responsible for impaired locomotion remain to be determined, the resemblance between Y17E and UNC-34-null animals suggests a profound loss of UNC-34 function. Thus, the importance of this conserved EVH1 residue extends from actin assembly in mammalian cells to neuronal migration and coordinated movement of an intact multicellular organism.

An important question arising from these findings concerns the kinase responsible for phosphorylation of Y16. Previous work identified Y39, but not Y16, as a major target of Abl1-dependent phosphorylation and implicated this modification in regulating VASP localization and interactions with zyxin ^41^. In agreement with these observations, the amino-acid sequence surrounding Y16 differs from the preferred substrate recognition motif of Abl family tyrosine kinases, which favors alanine at position +1 and proline at +3 ^70^. This suggests that Y16 may be phosphorylated by a distinct tyrosine kinase, potentially a member of the Src family, particularly given that Ena/VASP proteins have been shown to interact with the SH3 domains of several non-receptor tyrosine kinases, including Src, Fyn and Lyn, through their PRD ^39,43^. Its location within the surface-exposed aromatic triad would make this residue accessible to regulatory phosphorylation, although ligand binding might also influence its accessibility. Identifying the responsible kinase, the physiological conditions under which Y16 becomes phosphorylated and the signaling pathways controlling this modification will therefore be important objectives of future studies.

The potential significance of such a regulatory mechanism is underscored by previous pharmacological studies targeting the same EVH1 recognition interface. Competitive peptidomimetic inhibitors developed by Opitz and colleagues displaced VASP from FAs and lamellipodial tips and suppressed invasion of MDA-MB-231 breast cancer cells ^71^. Subsequent development of nanomolar-affinity inhibitors further demonstrated impaired tumor cell invasion and extravasation ^72^. The striking similarities between pharmacological blockade of EVH1 ligand-binding and the consequences of Y16E substitution suggest that phosphorylation of this residue could provide an endogenous means of modulating the same interactions. Although the physiological occurrence and regulation of this modification remain to be established, our findings identify the canonical EVH1 recognition surface as a potentially important point of convergence between intracellular signaling and spatial control of Ena/VASP activity.

## Methods

### Constructs

For the expression of proteins in *E. coli* the following constructs were generated. The coding sequence of mVASP was excised by BamHI-SalI from pGEX-6P-1-mVASP ^28^ and inserted into the same sites of pETDuet-1 (Novagen) to yield pETDuet-mVASP-WT. Phosphomimetic Y16E and Y39E variants of mVASP were generated by PCR-based site-directed mutagenesis using pETDuet-mVASP-WT as template, following standard procedures. As recombinant ActA is highly susceptible to proteolytic degradation in *E. coli*, a construct enabling tandem-affinity purification (TAP) was generated to facilitate purification to homogeneity. The internal region of *L. monocytogenes* EGD ActA harboring all four FP_4_ motifs (residues 248-389) was amplified from plasmid pSPL21 ^10^ by PCR using the primers ActA-BU 5’-TCTGGATCCTGGAAAAAGAAAAGTGAAGAGGTAAAT-3’ and ActA-SD 5’-GAGTTGAACGGGAGAGGCGGTAGATGGCATCACCATCACCATCACCATCACTGCTAGGTCGACTGT-3’. The resulting PCR product was inserted into the BamHI and SalI sites of pGEX-6P-1 (GE Healthcare), yielding pGEX-6P1-ActA-248-389-His_8_-C. Both primers additionally encoded a tryptophan residue to facilitate spectrophotometric detection of the ActA fragment by absorbance at 280 nm. Plasmid pGEX-6P1-SNAP for the expression of GST-SNAP was described previously ^21^.

For protein expression in mammalian cells, the following constructs were used: pmScarlet-LifeAct ^73^ as well as pEGFP-C1-mVASP-WT, pEGFP-C1-mEvl-WT, pEGFP-C1-mMena-WT ^28^. Phosphomimetic (Y16E, Y39E and S153D) and nonphosphorylatable (Y16F, Y39F and 153A) variants of the latter were generated by PCR-based site-directed mutagenesis using the corresponding WT constructs as templates and following standard procedures. Plasmids pEGFP-C1-mVASP-WTΔGAB/ΔFAB and pEGFP-C1-mVASP-Y16EΔGAB/ΔFAB, lacking residues 218-236 and 256-273, were generated by two successive rounds of PCR-based site-directed mutagenesis using pEGFP-C1-mVASP-WT and pEGFP-C1-mVASP-Y16E as the respective templates.

As stable expression of VASP from the CMV promoter-driven pEGFP-C1 vector could not be achieved, presumably due to promoter silencing, a pEGFP-C1 derivative was generated to allow stable expression under the control of the murine EF1α promoter and selection with puromycin. Briefly, the SV40 terminator was PCR-amplified from pPUR (Clontech) and inserted into the SalI-Acc65I sites of pEGFP-C1, yielding pEGFP-C1-SV40T. Next, the 1.3 kbp puromycin resistance cassette was amplified from pPUR and cloned into the Acc65I-BamHI sites of pEGFP-C1-SV40T, yielding pEGFP-C1-Puro. Subsequently, the mVASP-WT coding sequence was amplified from pEGFP-C1-mVASP-WT ^28^ and inserted into the BspEI-XhoI sites of pEGFP-C1-Puro, yielding pEGFP-C1-mVASP-WT-Puro. The CMV promoter was then replaced with a 1.2 kbp fragment encompassing the murine EF1α promoter and intron A, amplified from B16-F1 genomic DNA, using the AseI and NheI restriction sites, yielding pEGFP-EF1a-mVASP-WT-Puro. The corresponding phosphomimetic (Y16E and Y39E) and nonphosphorylatable (Y16F and Y39F) variants were generated by PCR-based site-directed mutagenesis using pEGFP-EF1a-mVASP-WT-Puro as the template. All generated constructs were validated by sequencing.

### Cell culture and transfection

B16-F1 mouse melanoma cells (CRL-6323) were obtained from ATCC. Ena/VASP-deficient triple-knockout mutants derived from B16-F1 cells and lacking Evl, VASP, and Mena (EVM-KO), together with mouse embryonic fibroblasts lacking all three Ena/VASP paralogues (MVE-KO) were described previously ^28^. B16-F1 WT and mutant cells were cultured at 37 °C and 5% CO_2_ in high-glucose DMEM culture medium (Lonza) supplemented with 10% FBS (Biowest), 2 mM UltraGlutamine (Lonza) and 1% Penicillin-Streptomycin (Biowest). B16-F1 and EVM-KO cells were transfected with 1 µg plasmid DNA and MVE-KO cells with 3 µg plasmid DNA, respectively using JetPRIME transfection reagent (PolyPlus) in 35 mm diameter wells (Sarstedt) according to the manufacturer’s instructions. Absence of mycoplasma in cell lines was routinely checked by the VenorGeM Mycoplasma Detection Kit (Sigma).

### Cultivation of *L. monocytogenes*

The wild-type strain of *L. monocytogenes* (EGD) was cultured at 37 °C under constant agitation in sterile brain heart infusion (BHI) broth (BD Biosciences). Infection experiments were performed using overnight cultures inoculated from single colonies.

### *Listeria* infection of B16-F1 cells

B16-F1 mouse melanoma cells were infected with *L. monocytogenes* at a multiplicity of infection (MOI) of 100 for 30 minutes at 37 °C in the presence of 7.5% CO2 in DMEM with 4.5 g/L glucose (Life Technologies) supplemented with 10% fetal calf serum (Capricorn Scientific) and 2 mM glutamine (Life Technologies). First, bacteria were harvested from an overnight culture by centrifugation at 3000 x g for 2 minutes and washed twice with DMEM without additives. Following MOI calculation and addition of the calculated volume of bacteria suspension to the cells, the infection process was initiated by centrifugation at 935 x g for 5 minutes. After a 30-minute infection period, cells were incubated in growth medium containing 50 µg/mL gentamicin (Sigma) for an additional 30 minutes to kill extracellular bacteria. Finally, cells were washed three times with DMEM before being overlaid with fresh growth medium containing 10 µg/mL gentamicin for further incubation. Subsequently, cells were processed for fixation and extraction or live-cell microscopy as detailed below.

### Fluorescence-Activated Cell Sorting

Cells stably expressing pEGFP-mVASP constructs were selected and maintained in growth medium supplemented with 2 µg/mL puromycin for at least two weeks. Subsequently, the cells were harvested, resuspended in MACS buffer containing 1 x PBS, pH 7.4 supplemented with 1 mg/mL bovine serum albumin, 1 mM EDTA, 25 mM Hepes and 0.1 mg/mL Normocin (Invivogen), and sorted into populations exhibiting low, intermediate, or high EGFP expression levels using fluorescence-activated cell sorting (FACS) (for more details see Supplementary Fig. 7). Cells with the lowest expression level were used for experiments. FACS was performed with an FACSAria III Fusion flow cytometer (BD Biosciences) driven by FACSDiva Vers. 8.0.1. software. EGFP was excited with a laser at 488 nm, and emission at 530 nm was detected with a band-pass filter. EGFP-positive cells were collected into sterile 5 mL PP tubes (Sarstedt) containing cell culture medium supplemented with 2 µg/mL puromycin and 25 mM HEPES.

### Protein purification

For expression of His-tagged mVASP variants, respective constructs were transformed into *E. coli* strain Rossetta 2 (Novagen) and protein expression was induced with 750 μM isopropyl-β-D-thiogalactoside (IPTG) at 21 °C for 16 h. The cells were then harvested and resuspended in lysis buffer containing 50 mM NaH₂PO₄/Na₂HPO₄ pH 8.0, 300 mM NaCl, 10 mM imidazole, 1 mM EDTA, 5 mM ß-mercaptoethanol, 2 mM benzamidine, 5% (v/v) glycerol, 0.1 mM AEBSF (AppliChem) and 2 U/mL Benzonase (Merck) and lysed by sonication. Fusion proteins were subsequently purified from bacterial extracts by immobilized metal affinity chromatography (IMAC) using cOmplete His-Tag Purification Resin (Roche) according to the manufacturer’s instructions. For expression of GST-ActA-248-389-His_8_-C and GST-SNAP, respective constructs were transformed into *E. coli* strain Rossetta 2 (Novagen) and protein expression was induced with 750 μM isopropyl-β-D-thiogalactoside (IPTG) at 21 °C for 16 h. The cells were then harvested and resuspended in lysis buffer containing 30 mM HEPES, pH 7.4, 150 mM NaCl, 1 mM EDTA, 2 mM benzamidine, 1 mM DTT, 5% (v/v) glycerol, 0.1 mM AEBSF (AppliChem) and 2 U/mL Benzonase (Merck) and lysed by sonication. Fusion proteins were subsequently purified from bacterial extracts by affinity chromatography using Protino Glutathione Agarose 4B (Macherey-Nagel) employing standard procedures. In case of GST-ActA-248-389-His_8_-C the GST-tag was cleaved with PreScission protease (GE Healthcare) and tandem affinity purification of the protein completed by IMAC purification using cOmplete His-Tag Purification Resin as stated above. In a final purification step, the proteins were further purified by size-exclusion chromatography (SEC) using HiLoad 26/600 Superdex 200 or HiLoad 26/75 Superdex (GE Healthcare) controlled by an Äkta Purifier System. Heterodimeric capping protein CapZ, profilin I (Pfn1) and SNAP-GCN4-Lpd^850–1250^(44A) were purified as previously described ^13,20^. Actin was isolated from rabbit skeletal muscle acetone powder using an established protocol ^74^. Fractions were labelled on Cys374 with ATTO488-maleimide (ATTO-TEC) and stored in G-Buffer (5 mM Tris/HCl, pH 8.0, 0.2 mM ATP, 0.5 mM DTT, 0.2 mM CaCl_2_, 0.1 mg/mL NaN_3_). GST-SNAP was labelled on cysteine residues with ATTO647-maleimide (ATTO-TEC). Protein concentrations were determined by absorption spectroscopy using extinction coefficients predicted from the amino-acid sequences.

### Analytical ultracentrifugation (AUC)

Sedimentation velocity runs were carried out in an analytical ultracentrifuge Optima AUC (Beckman Coulter, USA) with an An-50 Ti rotor at 50,000 rpm and 10 °C. Concentration profiles were measured with the absorption scanning optics at 280 nm using 12 mm standard double-sector centerpieces filled with 400 μL sample. Experiments were performed in a buffer containing 137 mM NaCl, 2.7 mM KCl, 1.8 mM KH_2_PO_4_, 10 mM Na_2_HPO_4_, 3 mM β-MESH, 0.1 mM EDTA, 0.1 mM AEBSF, pH 7.4. To compare the interaction of mVASP-WT and the Y16E and Y39E mutants with ActA, 2.5 µM of each VASP_4_ protein (*M*=165.1 kDa) was titrated with an ActA fragment (residues 248-389, containing all four FP_4_ motifs, *M*=17.7 kDa), using molar ratios of 1:0.5, 1:1, 1:2, 1:4 and 1:8. After mixing, samples were equilibrated for 4 h at 10 °C before sedimentation analysis.

For data analysis, a model for diffusion-deconvoluted differential sedimentation coefficient distributions (continuous *c*(*s*) distributions) implemented in the program SEDFIT ^75^ was used. Protein concentrations were determined spectrophotometrically, and extinction coefficients at 280 nm calculated from amino acid composition using the program SEDNTERP 3 ^76^. Throughout the text, concentrations of ActA are given in monomer units, and those of His-mVASP-WT and the Y16E and Y39E mutants in tetramer units. Integration of the *c*(*s*) distributions to quantify free ActA and the generation of AUC figures were performed with the program GUSSI ^77^.

### Preparation of small unilamellar vesicles

Small unilamellar vesicles (SUVs) were generated from 1,2-dioleoyl-sn-glycero-3-phosphocholine (DOPC; Avanti Polar Lipids) using a Mini-Extruder. Briefly, chloroform was removed from the DOPC stock solution by evaporation under a dry nitrogen stream for 1 h. The resulting lipid film was rehydrated to a final concentration of 2 mg/mL in vesicle buffer containing 20 mM HEPES pH 7.4, 150 mM NaCl, 5 mM KCl, 3 mM MgCl₂ and 5 mM β-mercaptoethanol. Following sonication for 3 min using a tapered microtip, the lipid suspension was passed 15 times through a polycarbonate membrane with 100 nm pores (Avanti Polar Lipids) to produce SUVs. SUVs were stored on ice and used within 5 days.

### *In vitro* TIRF microscopy

For TIRF imaging, glass coverslips were thoroughly cleaned by successive 15 min incubations in 2% (v/v) Hellmanex III dissolved in bidistilled water (ddH₂O) at 50 °C, 1 M KOH, 1 M HCl and 70% ethanol, with extensive ddH₂O washes between each step. Following air-drying, coverslips were treated for 30 min with ambient air plasma in a benchtop plasma cleaner (Harrick Plasma). Two-channel flow chambers were then constructed by attaching the cleaned coverslips to glass slides using three narrow strips of double-sided adhesive tape as spacers. Supported lipid bilayers were generated by introducing 20 µL of the SUV suspension into each channel and allowing the vesicles to spread across the glass surface for 30 min. Unbound material was subsequently removed by washing twice with 1× KMEI buffer containing 1% BSA. For TIRF bead assays, 50 μL of SNAP-capture magnetic beads with a diameter of 2 μm (NEB #S9145Z) were washed twice with coupling buffer (20 mM HEPES pH 7.4, 150 mM NaCl, 2 mM DTT, 2 mM β-mercaptoethanol, 10% BSA) and then coated with 10 μM of SNAP-GCN4-Lpd^850–1250^(44A) for 16 h on a wheel rotator at 4 °C to obtain lamellipodin-derivatized beads. Subsequently, the beads were washed three times with coupling buffer and then incubated with 0.5 μM soluble His-mVASP-WT, His-mVASP-Y16E or His-mVASP-Y39E in coupling buffer for 1 h at 4 °C on a wheel rotator. Subsequently, beads were washed five times with coupling buffer to remove unbound VASP and then re-suspended into storage buffer (20 mM HEPES pH 7.4, 150 mM NaCl, 2 mM DTT, 2 mM β-mercaptoethanol, 10% BSA and 60% glycerol) for direct use in experiments or storage at -20 °C. TIRF assays were performed in TIRF buffer (20 mM imidazole pH 7.4, 1 mM MgCl_2_, 50 mM KCl, 0.5 mM ATP, 1 mM EGTA, 20 mM β-mercaptoethanol, 15 mM glucose, 20 μg/mL catalase, 100 μg/mL glucose oxidase, and 2.5 mg/mL methylcellulose (4,000 cP)). Actin was polymerized together with 5 µM Pfn1 in the presence of 25 nM CP and coated beads. Actin assembly was initiated by addition of ATP-G-actin (3 μM, 10% Atto488-labelled), followed by immediate transfer of the reaction mixture into the prepared flow chambers. Images were captured at 5 s intervals for at least 20 min using a Nikon Eclipse Ti-E inverted microscope equipped with a CFI Apochromat TIRF 100×/1.49 NA oil-immersion objective and iXon3 897 back-illuminated EMCCD cameras (Andor), with 488 nm and 647 nm laser excitation and exposure times of 70 ms.

### Antibodies

For immunoblotting, polyclonal antibodies directed against mVASP (1:1,000) ^28^, or mouse monoclonal antibodies against glyceraldehyde-3-phosphate dehydrogenase (GAPDH) (1:10,000; #CB1001-500UG, Merck-Millipore) were used. Primary antibodies in immunoblots were visualized using phosphatase-coupled anti-rabbit (1:1,000; #115-055-144, Dianova) or anti-mouse antibodies (1:1,000; #115-055-62, Dianova). For immunofluorescence, the following primary antibodies were used: polyclonal rabbit anti-VASP antibody (1:1,000 dilution ^28^), mouse monoclonal anti-vinculin antibody (1:1,000 dilution; #V9131, clone hVIN-1, Sigma) and polyclonal anti-*Listeria* antibody (1:100 dilution, K52 ^63^). Primary antibodies were visualized in immunocytochemistry with Alexa-350-conjugated goat anti-rabbit IgG (1:200 dilution; #A11046, Invitrogen), Alexa-488-conjugated goat-anti-rabbit (1:1,000 dilution; #A-11034, Invitrogen), Alexa-633-conjugated goat-anti-rabbit (1:1,000 dilution; #A-21070, Invitrogen) or Alexa-555-conjugated goat-anti-mouse (1:1,000 dilution; #A32727; Invitrogen) antibodies.

### Immunofluorescence

For immunofluorescence labeling, cells were seeded on glass coverslips (ibidi) coated with either laminin (25 μg/mL) or fibronectin (10 μg/mL) and allowed to adhere for 2 h. Cells were then fixed for 20 min in pre-warmed PBS pH 7.4 containing 4% paraformaldehyde (PFA) and 0.06% picric acid, followed by three washes with PBS containing 100 mM glycine. After permeabilization with 0.05% Triton X-100 in PBS for 30 s, specimens were blocked in PBG (PBS containing 0.5% BSA) for 30 min and subsequently incubated with primary antibodies diluted in PBG overnight at room temperature. Following three washes with PBG, secondary antibodies were applied for at least 2 h. EGFP fluorescence was enhanced with an Alexa488-conjugated anti-GFP nanobody ^78^, whereas filamentous actin was visualized with ATTO550-phalloidin (ATTO-TEC; #AD550-8, 1:250). Samples were mounted in Mowiol (Sigma) and images captured using either an inverted Olympus IX-81 microscope equipped with a UPlan FI 100x/1.30 NA oil-immersion objective or a Zeiss LSM 980 confocal microscope with Airyscan 2 and a Plan-Apochromat 63x/1.4 NA oil DIC objective. Images were processed and analyzed using Fiji software ^79^. Fluorescence intensity profiles from line scans of antibody-stained *L. monocytogenes* were measured in images acquired under identical imaging settings. To quantify FA localization of EGFP-tagged mVASP variants, colocalization with vinculin was assessed in maximum-intensity projections of confocal z-stacks generated in Fiji. Pearson’s correlation coefficients (*r*) were subsequently calculated using the Coloc2 plugin within regions of interest excluding the leading edge.

### Quantification of *Listeria* actin-tail length

To quantify the length of *L. monocytogenes* actin tails, B16-F1 cells were seeded onto 12-mm ethanol/acid-washed glass coverslips pre-coated for 1 h at room temperature with 25 µg/mL laminin (Sigma) in coating buffer (50 mM Tris-HCl pH 7.4, 150 mM NaCl). The following day, cells were infected with wild-type *L. monocytogenes* as described above. To allow actin tails to develop, infected cells were incubated for an additional 5 h in growth medium containing 10 µg/mL gentamicin before fixation with 4% pre-warmed paraformaldehyde (PFA) in PBS for 20 min. Following fixation, cells were washed twice with PBS and permeabilized for 1 min with 0.05% Triton X-100 in PBS, followed by blocking for 1 h at room temperature with 5% horse serum in 1% BSA/PBS. Primary antibodies were diluted in 1% BSA/PBS and incubated with the samples for 1 h at room temperature. *L. monocytogenes* was stained using the polyclonal primary antibody K52.

After three washes with PBS, samples were incubated with secondary antibodies diluted in 1% BSA/PBS according to manufacturers’ instructions for 1 h at room temperature. Alexa-350-conjugated goat anti-rabbit IgG (Invitrogen) was then used to visualize *L. monocytogenes*. ATTO594-labelled phalloidin (ATTO-TEC; #AD-594-81, 1:200) was added to the secondary antibody solution to stain F-actin. EGFP fluorescence was routinely enhanced using ATTO488-labeled anti-GFP nanobodies ^28^ at a dilution of 1:1,000. Finally, coverslips were washed three times with PBS, mounted using ProLong™ Diamond Antifade Mountant (Invitrogen), and stored at 4 °C until image acquisition.

Imaging was performed using a CSU-W1 spinning disk confocal module (Yokogawa Electric) and a Zyla 4.2 sCMOS camera (Andor) mounted on a Nikon Eclipse Ti2 microscope (Nikon). Images were acquired using a Plan Fluor 100x/1.3 NA oil-immersion objective (Nikon), 405/488/561/638 nm laser lines (Omicron), and 447/60, 525/50 as well as 600/52 single-band emission filters, all controlled by NIS-Elements software (Nikon). Z-stacks were acquired using a piezoelectric z-drive (Mad City Labs) with a step size of 200 nm. Actin tail lengths were quantified from maximum-intensity projections of the acquired z-stacks using the Manual Tracking plugin in Fiji/ImageJ. To avoid overrepresentation of heavily infected cells, a maximum of three actin tails per cell were measured, with the three longest tails selected for quantification in each condition.

### Live cell imaging

Time-lapse imaging of cells was performed using an Olympus XI-81 inverted microscope (Olympus) controlled by Metamorph software (Molecular Devices) and equipped with objectives specified below and a CoolSnap EZ camera (Photometrics). Cells were seeded onto 35 mm glass bottom dishes (ibidi) coated with 25 μg/mL laminin (Sigma), and maintained in imaging medium composed of F-12 Ham Nutrient Mixture with 25 mM HEPES (Sigma) to compensate for the lack of CO_2_, and supplemented with 10% FBS (Biowest), 1% Penicillin-Streptomycin (Biowest), 2 mM stable L-glutamine (Biowest), and 2.7 g/mL D-glucose (Carl Roth) in an ibidi stage top heating system at 37 °C. For random motility assays, cells were seeded at low density onto dishes and allowed to adhere for 3 h. Subsequently, the growth medium was exchanged with imaging medium, the chamber mounted into the heating system, and cells recorded by phase-contrast, time-lapse imaging at 1 min intervals for 3 h using an Uplan FL N 4x/0.13NA objective (Olympus) with 1.6x Optovar magnification. Single cell tracking was performed in ImageJ byMTrackJ ^80^. Analyses of cell speed and mean square displacements were performed in Excel (Microsoft) using a customized macro ^81^. Cells that contacted each other or divided were excluded from analyses. Lamellipodial persistence of randomly migrating cells on laminin was determined from phase-contrast, time-lapse movies. Lamellipodial persistence was defined as time in min from initiation until collapse of the lamellipodium. To visualize VASP and actin dynamics during cell migration, cells stably expressing EGFP-mVASP-WT or the corresponding phosphomimetic and nonphosphorylatable variants were transfected with pmScarlet-LifeAct. At 16 h post-transfection, cells were seeded onto laminin-coated (25 μg/mL) glass-bottom dishes (ibidi) and allowed to adhere and spread for 1 h in imaging medium supplemented with 0.1 mM Trolox (Sigma). Migrating cells were subsequently imaged by multicolor time-lapse epifluorescence microscopy using an Olympus IX83 inverted microscope equipped with an UPlanApo 60x/1.50 Oil HR TIRF objective, an ORCA-Flash4.0 camera, and a temperature-controlled incubation system set to 37 °C. Images were acquired at 10 s intervals for at least 10 min.

### Analysis of bacterial movement speed

To quantify the intracellular movement speed of *L. monocytogenes*, high-magnification time-lapse phase-contrast and fluorescence microscopy was performed. For this purpose, B16-F1 WT or EVM-KO cells transfected with different EGFP-mVASP variants were seeded onto laminin-coated glass-bottom dishes (ibidi) and infected as described above. For illustrative movies 3 and 4 not used for quantification, cells were additionally co-transfected with mScarlet3-actin to avoid potential effects of exogenous actin expression on actin tail formation rates in quantitative analyses. After incubation for 3 h in growth medium containing 10 µg/mL gentamicin, cells were washed three times with pre-warmed, phenol red-free, HEPES-buffered Ham’s F-12 Nutrient mixture medium (Sigma) supplemented with 10% FCS (Capricorn), 2 mM glutamine, and 10 µg/mL gentamicin. Following medium exchange, dishes were secured in a microscope heating chamber (PeCon) maintained at 37 °C and mounted on an Axio Observer.A1 microscope (Carl Zeiss) equipped with a VIS-LED illumination system for phase-contrast imaging, a DG4/30 xenon arc lamp (Sutter Instrument), a dual FITC/mCherry filter set, and a Plan-NEOFLUAR 100×/1.3 NA Ph3 oil-immersion objective (Carl Zeiss). Images were acquired every 5 s for quantitative analyses or every 10 s for illustrative movies over a period of 5-10 min using a pco.edge 4.2 LT sCMOS camera (Excelitas Technologies) controlled by VisiView® software (Visitron Systems).

Bacterial movement speeds were determined using the Manual Tracking plugin in Fiji/ImageJ by tracking the rear pole of each bacterium adjacent to the actin tail and calculating the distance traveled over time. To avoid overrepresentation of heavily infected cells, a maximum of three bacteria per cell were analyzed. Given the high variability in bacterial movement speeds and their potential dependence on host cell size and bacterial load, the three fastest bacteria moving freely within the host cytoplasm for at least 20 s were selected for quantitative comparisons.

### *C. elegans* strains and handling

*C. elegans* strains used in this study: N2 - wild type; CB566 - *unc-34(e566)* V; RZB500 - *unc-34 (msn214[Y17E])* V; RZB525 - *unc-34 (msn216[Y17F])* V. All strains were grown and maintained on nematode growth medium (NGM) plates seeded with OP50 bacteria according to standard protocols ^66^. All worms were kept at 20 °C.

### Generating the *unc-34* mutant strains

CRISPR-Cas9-mediated genome editing was used to generate *unc-34* phospho-mutant worms ^82^. An sgRNA was selected with a PAM site located 10 bp from the codon encoding tyrosine 17 in the unc-34 gene, targeting the antisense strand: 5’-ATTCGTTGTAGGTCATCACT-3’. The HDR repair templates (single stranded oligodeoxynucleotide; ssODN) were designed with multiple point mutations to change amino acid 17 from Y to either E or F, as well as silent mutations to prevent reattachment and recutting by the Cas9 machinery. The short, mutated sequences were flanked by 35 bases long homology arms. The introduction of silent mutations created a new restriction site for the BspHI restriction enzyme, which facilitated genotyping by PCR.

Y17E ssODN: AGCGAAGCATGTATCGCGACGGCAATGGCCCGAGTCATGACA**GAA**AATGAGTCG ACAAAAGGATGGGTACTACTCGGCGGAAATGA

Y17F ssODN: AGCGAAGCATGTATCGCGACGGCAATGGCCCGAGTCATGACA**TTT**AATGAGTCG ACAAAAGGATGGGTACTACTCGGCGGAAATGA

Gonads of one day old adult N2 worms were injected with the following mix: Cas9 enzyme (0.8 μg/μL), *unc-34* sgRNA (0.06 mM) and ssODN (0.225 μg/μL), *dpy-10* sgRNA (0.02 μg/μL) and *dpy-10* ssODN (0.04 μg/μL), KCl (25 mM) and tracrRNA (0.1 μg/μL). The F1 progeny were screened phenotypically for dumpy worms. F1 dumpy worms were isolated, allowed to lay embryos, and later screened for the *unc-34* mutation by PCR using the following primers: forward primer: 5’-TTTCGATTTCCGGCCACCG-3’ reverse primer: 5’-TGTCGCATCCTGTTCATTCTCG-3’, and by cutting the PCR product with BspHI (New England Biolabs). All CRISPR reagents and primers were purchased from IDT (Integrated DNA Technologies). Homozygous mutants were confirmed by sequencing and subsequently outcrossed to eliminate the dumpy phenotype and minimize potential off-target mutations.

### High-temporal-resolution imaging of L4-stage worm movement

To capture worm locomotion at high temporal resolution over a large field of view, synchronized L4-stage worms were transferred with an eyelash onto 60 × 15 mm NGM plates containing freshly seeded bacterial lawns. Brightfield time-lapse movies were acquired at 10x magnification using a Nikon SMZ18 stereomicroscope equipped with an SHR Plan Apo 1x WD 60 objective, a Nikon DS-Fi3 high-definition color camera with a 5.9-megapixel CMOS image sensor, and a Nikon C-0.55x DS C-mount adapter. Each movie was recorded for 20 s at 15 frames per second, with a spatial resolution of 679 pixels/mm and a frame size of 2880 x 2048 pixels. These recordings were subsequently used to generate the temporally color-coded projections shown in Fig. 6c. All image processing and analyses were performed using Fiji/ImageJ. Briefly, 20 s time-lapse recordings were reduced to 150 frames, and individual worms were segmented by manually adjusting and applying an intensity threshold. Gaps in the segmented worm outlines were closed using the “Close-” and “Fill Holes” functions. Subsequently, worm regions of interest (ROIs) were generated for each frame using “Analyze Particles,” and their outlines were drawn with a line width of 2 pixels. Finally, temporally color-coded projections of the worm outlines were generated using the “Temporal-Color Code” function with the Fire lookup table (LUT) to visualize their positions over time.

### Multi-worm tracking

To analyze worm locomotion, 10-15 late L4 worms were transferred to a single 60 mm NGM plate with a freshly seeded bacterial lawn. Each plate was recorded for 2 minutes at 2 frames per second with a resolution of 103 pixels/mm in the worm tracker machine. The tracker machine was custom-built by commercial sources affiliated with Behavioral Phenomics Lab of André Brown (MRC Laboratory Institute of Medical Sciences). The tracker is fitted with a Genie Nano M2420 Mono camera (Teledyne Dalsa) on top and red-light source from below. The acquired videos underwent image analysis for feature extraction by Tierpsy software ^83^. In this software, the raw video files were compressed and processed using an adaptive thresholding technique that subtracts the background, creating a mask around the body of the worm to predict the worm backbone. For all worms in the movie the software draws trajectory lines to differentiate individual worm paths. Worm trajectories of colliding worms or worms moving out of the frame were excluded from the analysis.

Worm locomotion was quantified using the ‘Speed’ function in Tierpsy, which measures whole-body movement in µm/s, with positive, zero and negative values indicating forward movement, pauses and backward movement, respectively. Median speeds across all worms in each video were plotted over time. Worm body curvature was quantified using the ‘Curvature_Std_Midbody’ function in Tierpsy, which calculates the standard deviation (SD) of local curvature along the midbody segment at 0.5-s intervals. Curvature was defined as the change in tangent angle along the body midline per unit length, with low SD values indicating smooth, uniform bending and high values reflecting uncoordinated and irregular body postures.

### Enhanced sampling umbrella MD simulations

The models chosen for this work are based on the crystal structure of the murine Mena-EVH1 domain in complex with the consensus proline-rich motif FPPPP (ACTA) peptide (PDB: 1EVH) ^53^. For calculations, the models of EVH1-WT, EVH1-Y16E and EVH1-pY16 were examined. The Y16E mutation was generated from the WT using the „Simple mutate” tool in Coot v0.9.8.95 ^84^. Phosphorylation of Y16 (pY16) was included by the „Build Structure/Modify Structure” tool in UCSF Chimera v1.16 ^85,86^, developed by the Resource for Biocomputing, Visualization, and Informatics at the University of California, San Francisco, with support from NIH P41-GM103311. All calculations were performed using the GROMACS package v2024.3 ^87^ applying parameters from the AMBER99SB force field ^88,89^ to all species in the simulated systems with six independent MD runs for each complex. For pY16, the force field implemented in GROMACS was supplemented with parameters for phosphorylated tyrosine ^90^. The umbrella sampling simulations were performed on the Halime high-performance computational cluster at the Institute for Biophysical Chemistry at Hannover Medical School, essentially as described previously ^91^. The generated models were used as starting configurations for the pulling simulations. The complexes were placed in rectangular simulation boxes of 5 × 5 × 10 Å, sufficiently large to satisfy the minimum image convention and accommodate pulling simulations along the z-axis. The TIP3P three-point rigid water model ^92^ was used and 150 mM NaCl was added, including neutralizing counterions. Calculations were run at 2 fs integration steps. Short-range non-bonded interactions were cut off at 1.4 nm and long-range electrostatics were treated with the particle mesh Ewald (PME) method ^93,94^ under periodic boundary conditions. Following steepest descents minimization, each of the systems underwent constant pressure equilibration (NPT) at 1.0 bar for 500 ps using the Berendsen weak coupling method ^95^ applying harmonic position restraints. Following equilibration, restraints were removed from the ActA peptide leaving EVH1 as immobile reference for the pulling simulations. During each simulation, the ActA peptide was pulled away from the EVH1 structure along the z-axis over 150 ps at a pull rate of 0.01nm ps^-1^, applying a spring constant of 1000kJ mol^-1^ nm^-2^ up to final center-of-mass (COM) distances between EVH1 and ActA of approximately 3 nm. These trajectories were used to generate the starting configurations for the umbrella sampling windows. An asymmetric distribution of sampling windows was used, such that the window spacing was 0.05 nm up to 2.5 nm COM separation, and 0.2 nm between 2.4 nm and 3 nm of COM separation, resulting in around 25 windows. In each window, 25 ns of productive MD was calculated for a total simulation time of approximately 625 ns. The spring constants were set to 1000 kJ mol^-1^ nm^-2^ or 5000 kJ mol^-1^ nm^-2^ for the 0.05 nm or 0.2 nm windows, respectively. The Nosé-Hoover thermostat ^96,97^ was used to maintain temperature, and the Parrinello-Rahman barostat ^98,99^ was used to isotropically regulate pressure. Analysis of results was performed with the weighted histogram analysis method (WHAM) ^100^ by bootstrapping with 1000 resamples. The resulting potential of mean force (PMF) profiles were exported to OriginPro v2024 (OriginLab), where the curves were averaged using the „Average/Merge Multiple Curves” tool. The dissociation energies were calculated as the differences between the maxima and minima of the respective profiles. Molecular graphics in Fig. 2a and Fig. 4a were designed with PyMOL 3.1 or UCSF Chimera v1.16, respectively.

### Quantification and statistical analysis

Quantitative experiments were performed in triplicates. The number of independent biological replicates as well as the number of events or cells quantified for analysis is specified in respective figure legends. Raw data were processed in Microsoft Excel. No samples or data points were excluded from analysis. Statistical analyses were performed with Origin 2021 (OriginLab). Data normality was assessed using the Shapiro-Wilk test. Comparisons between two groups were performed using Student’s *t*-test for normally distributed data or the nonparametric Mann-Whitney *U* test for non-normally distributed data. For comparisons involving more than two groups, normally distributed data were analyzed by one-way analysis of variance (ANOVA) followed by Tukey’s multiple comparison test, whereas non-normally distributed data were analyzed using the Kruskal-Wallis test followed by Dunn’s multiple comparison test. Statistical differences were defined as \**P* < 0.05, \*\**P* < 0.01, \*\*\**P* < 0.001 as well as n.s., not significant, and are displayed and stated in figures and figure legends, respectively. Values are shown as box plots or as means ± SD or means ± SEM.

## Data availability

All data needed to evaluate the conclusions in the paper are present in the paper and/or the Supplementary Materials. Materials can be provided by J.F. pending scientific review and a completed material transfer agreement. Requests for the materials should be submitted to J.F.

## Acknowledgements

We thank Sandra Witting and Annette Breskott for excellent technical assistance and Matthias Ballmaier in the flow cytometry core unit of the Hannover Medical School for performing the cell sorting experiments.

## Author contributions

J.S., T.K., P.K., J.K., A.N., J.E., U.C., K.R., R.Z.-B. and J.F. performed experiments, interpreted data and/or made and edited Figures. J.S., J.F., K.R, and R.Z.-B. developed the project. J.S. and J.F. wrote large parts of the paper with conceptual help from K.R and R.Z.-B. All authors contributed to text corrections and proofreading.

## Funding

This work was supported by the Deutsche Forschungsgemeinschaft (DFG, German Research Foundation), grant Fa330/16-1 (to J.F.) and instrumentation grant INST 192/534-1 FUGG for the Beckman Coulter Optima AUC, intramural funding from the Helmholtz Association (to K.R.) and the Israel Science Foundation, grant 1677/25 (to J.E. and R.Z.B.).

## Competing interests

Authors declare that no competing interests exist

## Supplementary Information

## Supplementary Figures

**Supplementary Fig. 1:**
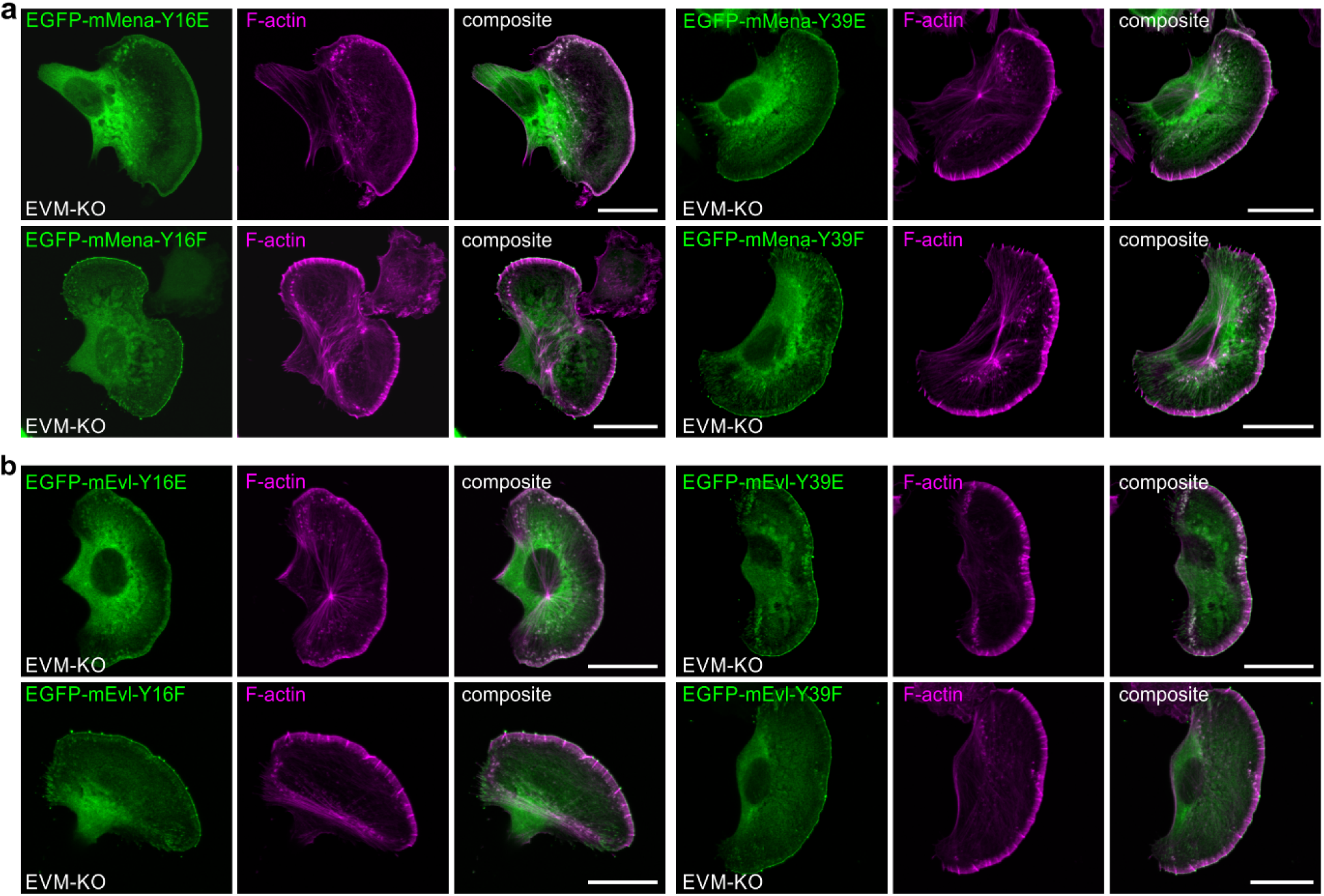
Phosphorylation of Y16, but not of Y39, blocks accumulation of mMena and mEvl at the tips of lamellipodia and microspikes. **a-b** Representative images of EVM-KO cells transiently expressing EGFP-tagged phosphomimetic or nonphosphorylatable mMena (**a**) and mEvl-variants (**b**) as indicated while migrating on laminin. Cells were fixed and stained with a GFP nanobody and phalloidin for the actin cytoskeleton. Brightest point projections of confocal stacks are shown. Scale bar, 20 μm. Note that microspike formation is not rescued in EVM-KO cells expressing EGFP-mMena-Y16E or EGFP-mEvl-Y16E, in contrast to cells expressing all other variants.

**Supplementary Fig. 2:**
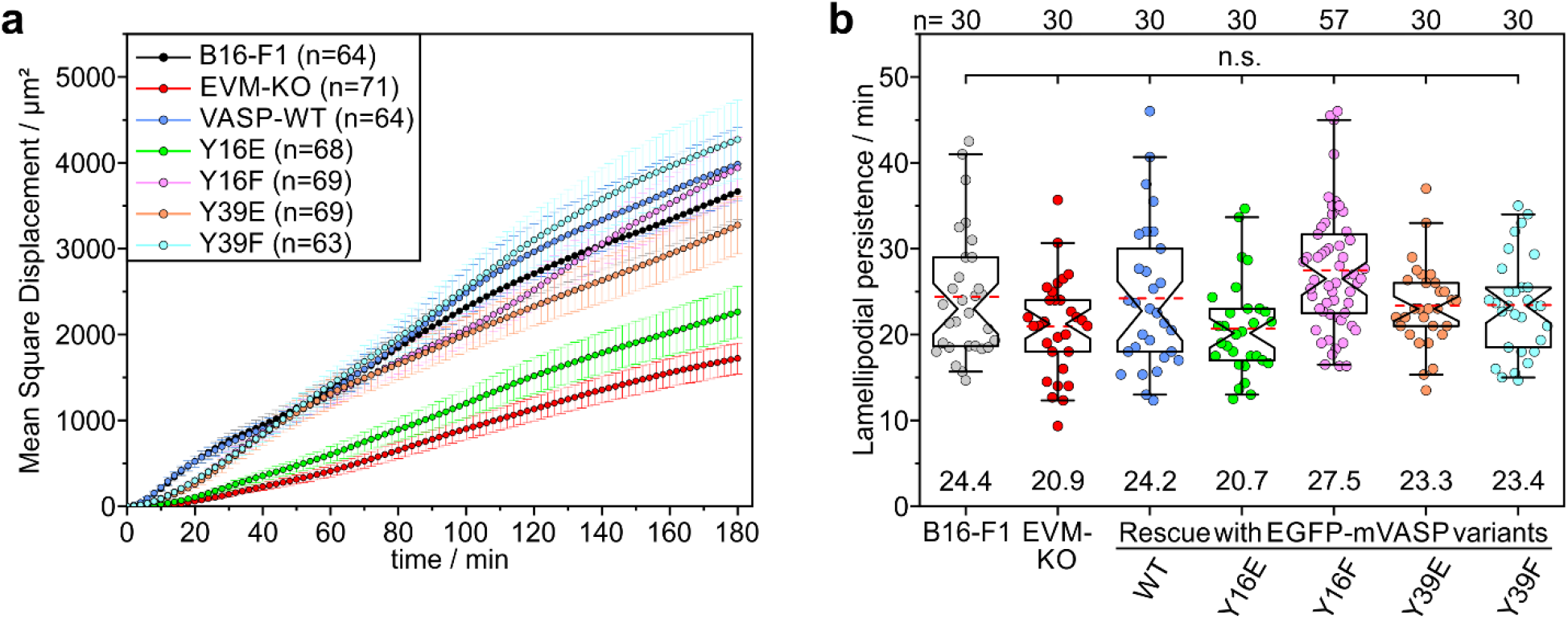
Phosphomimetic mVASP-Y16E fails to rescue EVM-KO cell migration. **a** Quantification of MSD in B16-F1 control cells, EVM-KO cells, and EVM-KO cell lines stably expressing the indicated EGFP-mVASP variants during migration on laminin. Data represent means ± SEM. Note that expression of phosphomimetic mVASP-Y16E in EVM-KO cells does not rescue their migration defect. **b** Quantification of lamellipodial persistence in the cell lines described above during migration on laminin. Although lamellipodial persistence was slightly increased in EVM-KO cells expressing nonphosphorylatable mVASP-Y16F, this difference was not statistically significant. Boxes in box plots indicate 50% (25 to 75%) and whiskers 90% (5 to 95%) of all measurements, with dashed red lines depicting the means. Medians are highlighted by indentation of boxes. Kruskal-Wallis test with Dunn’s multiple comparison test. n.s., not significant. (**a,b**) *n*, number of cells analyzed. Results are pooled data from at least three biologically independent replicates.

**Supplementary Fig. 3:**
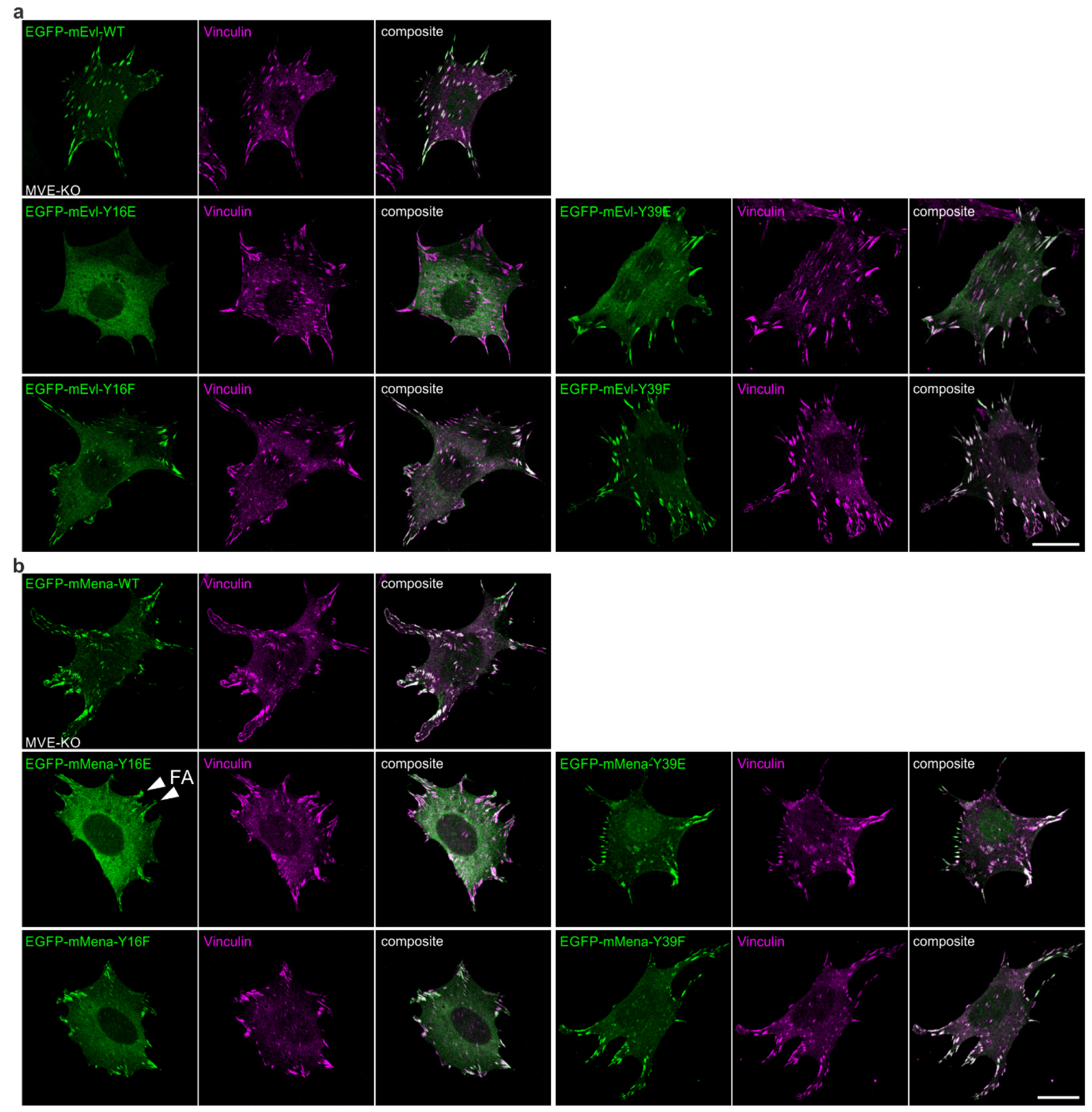
Mena recruitment to focal adhesions persists despite Y16E-mediated disruption of canonical targeting. **a-b** Representative images of Ena/VASP-deficient MVE-KO fibroblasts transiently expressing EGFP-tagged mEvl (**a**) or mMena (**b**) variants as indicated. Note that the Y16E substitution abolished mEvl targeting to FAs, whereas mMena-Y16E retained moderate FA localization (white arrow heads), presumably through its LERER repeats, which have been shown to mediate direct binding to α5 integrin ^58^. Brightest point projections of confocal stacks are shown. Scale bars, 20 μm.

**Supplementary Fig. 4:**
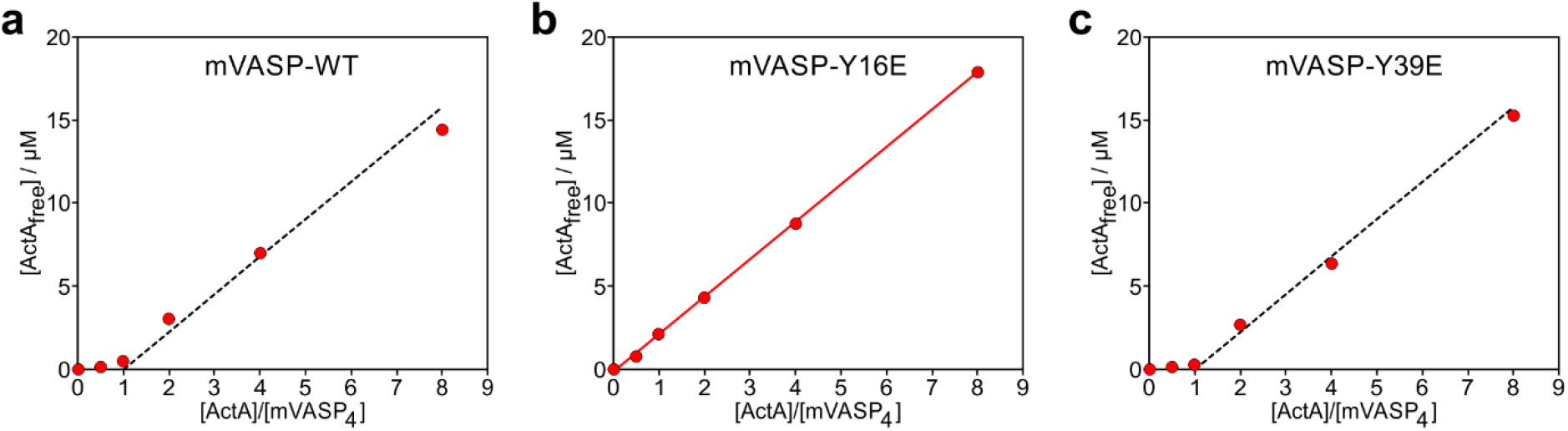
The mVASP tetramer forms an equimolar complex with ActA. **a-c** Free ActA concentrations were approximated from AUC experiments by integrating the 1 S peak in the *c*(*s*) distributions from titrations of mVASP_4_-WT (**a**), mVASP_4_-Y16E (**b**) or mVASP_4_-Y39E (**c**) with ActA at molar ratios of 0:1, 0.5:1, 1:1, 2:1, 4:1 and 8:1, with three representative concentrations shown in Fig. 4d-f. For VASP_4_-Y16E, which did not bind ActA, the amount of free ActA increased linearly with the molar ratio. In contrast, virtually no free ActA was observed for mVASP_4_-WT and mVASP_4_-Y39E up to a 1:1 molar ratio. For comparison, starting at a 1:1 molar ratio, lines with the slope of the linear regression for mVASP_4_-Y16E (red line in **b**) were overlaid as dashed black lines in **a** and **c**.

**Supplementary Fig. 5:**
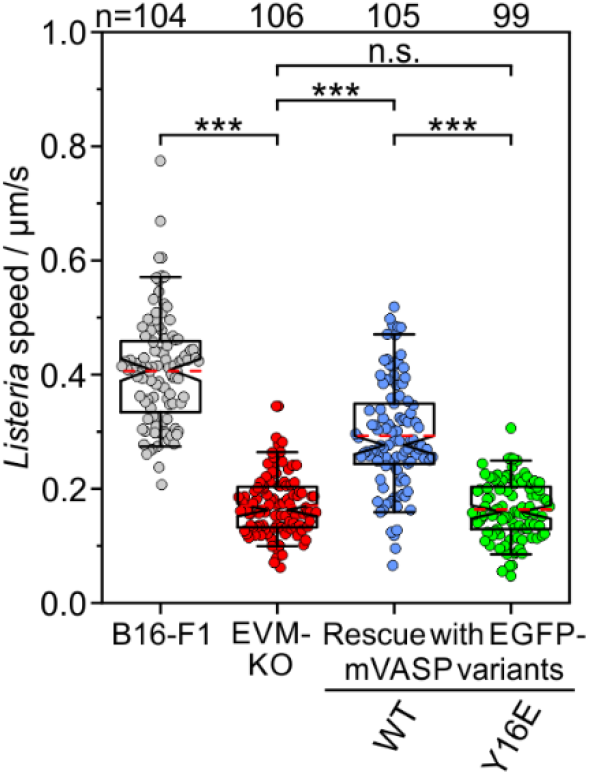
Phosphomimetic mVASP-Y16E fails to rescue intracellular *Listeria* motility. Quantification of *L. monocytogenes* motility in infected cells as indicated. Boxes in box plots indicate 50% (25 to 75%) and whiskers 90% (5 to 95%) of all measurements, with dashed red lines depicting the means. Medians are highlighted by indentation of boxes. Kruskal-Wallis test with Dunn’s multiple comparison test. *n*, number of bacteria analyzed. Results are pooled data from at least three biologically independent replicates. \*\*\**P* < 0.001. n.s., not significant.

**Supplementary Fig. 6:**
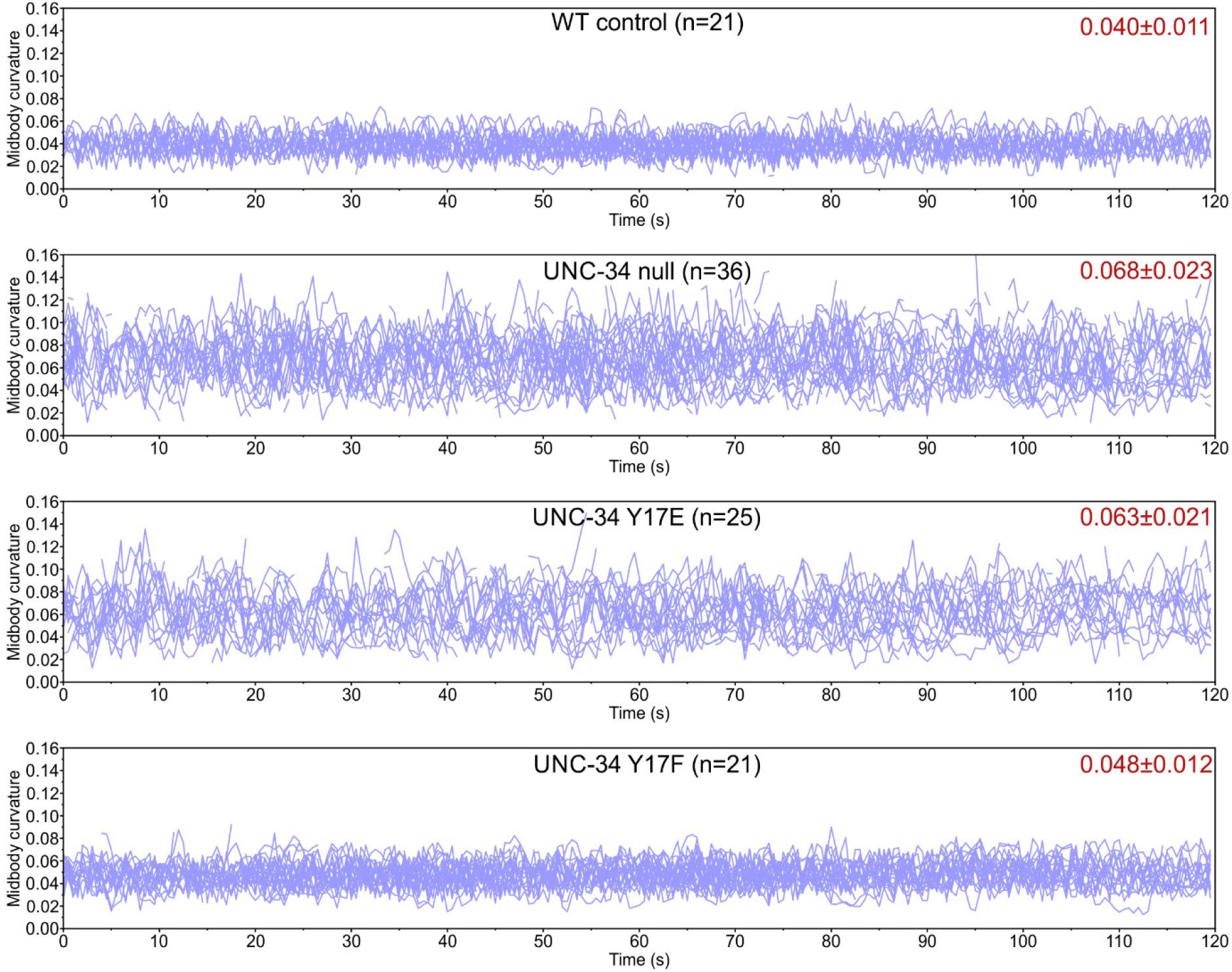
Phosphomimetic UNC-34-Y17E phenocopies worm curvature of UNC-34-null mutants. Quantification of midbody curvature over time in the indicated L4 larvae. Data points captured at 0.5 s intervals represent the standard deviation of curvature along the mid-body segment of each worm in the corresponding movie. Low values of mid-body curvature indicate smooth, uniform body contours, as observed in WT animals or those expressing nonphosphorylatable UNC-34-Y17F, whereas frequent sharp bends and high curvature values indicate uncoordinated, irregular body postures, as seen in UNC-34-null animals and those expressing phosphomimetic UNC-34-Y17E. Values represent mean ± SD across all time points. Each line represents the curvature values of a single worm. Results are pooled data from at least three biologically independent replicates. *n*, number of L4 worms.

**Supplementary Fig. 7:**
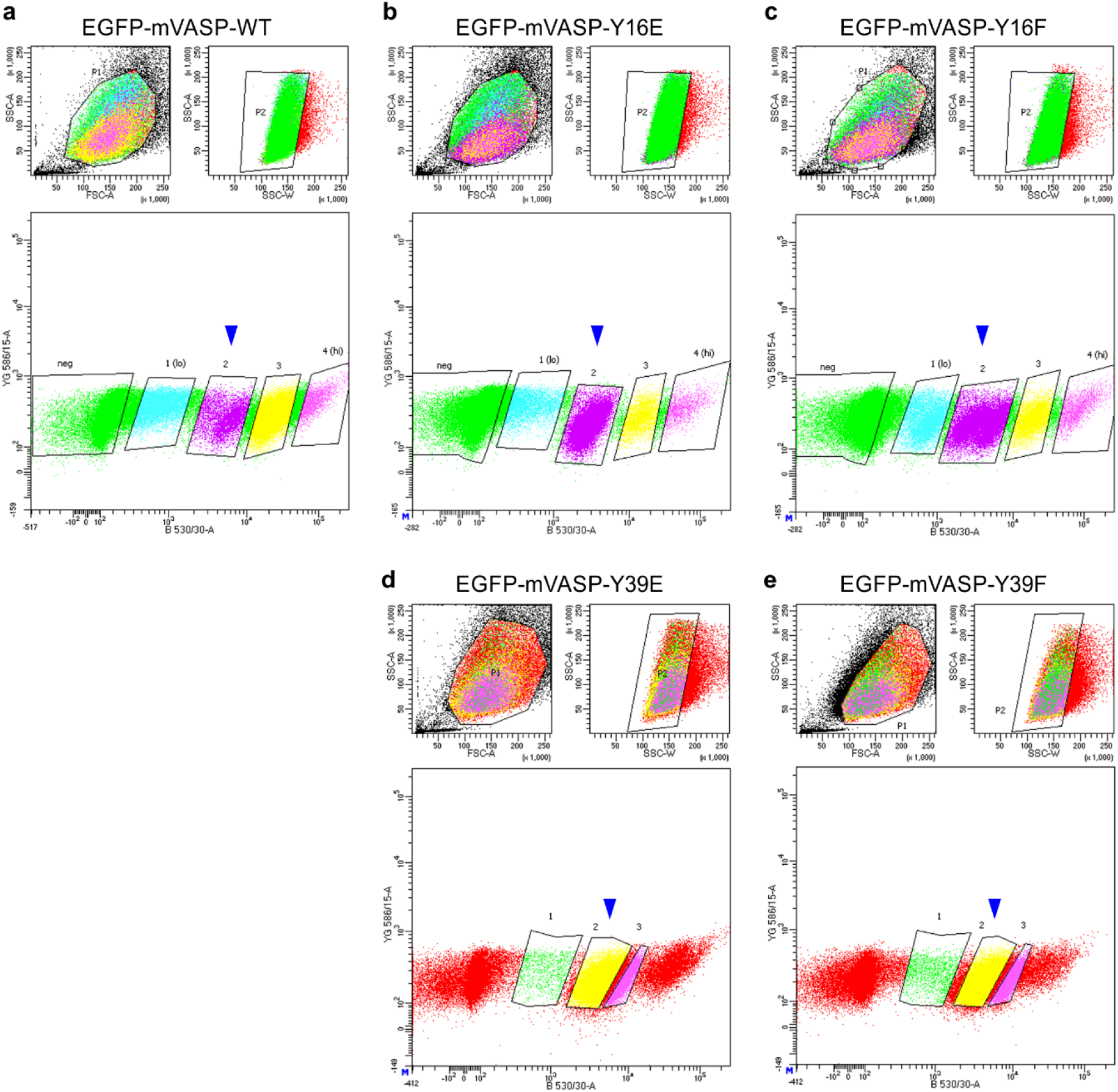
FACS data and gating strategy. **a-e** FACS plots of EVM-KO cells stably expressing EGFP-mVASP variants. *Upper left panels*: Gating of cells based on forward and side scatter (FSC-A *versus* SSC-A). *Upper right panels*: Exclusion of doublets using side scatter area and side scatter width (SSC-A *versus* SSC-W). *Bottom panels:* Gating of EGFP-positive cells (B 530/30-A versus YG 586/15-A) into populations with different expression levels, as indicated. Blue arrowheads indicate the low-expression populations selected for experiments. Each plot represents 100,000 recorded events per cell line.

## Movie Legends

**Movie 1: Phosphorylation of Y16, but not of Y39, blocks accumulation of VASP at the tips of lamellipodia and microspikes.** Representative epifluorescence, time-lapse movies of EVM-KO cells stably expressing EGFP-tagged phosphomimetic or nonphosphorylatable mVASP-variants as indicated together with the F-actin probe mScarlet-LifeAct while migrating on laminin. EVM-KO cells expressing mScarlet-LifeAct alone or additionally EGFP-mVASP-WT were used as controls. Note the lack of subcellular targeting to the tips of lamellipodia and microspikes as well as the failure to rescue microspike formation upon expression of phosphomimetic mVASP-Y16E, in contrast to the other mVASP variants. Time is indicated in min:sec. Scale bar, 20 µm.

**Movie 2**: **Surface-bound Lamellipodin does not recruit phosphomimetic mVASP-Y16E from solution to drive processive actin assembly.** Polymerization of 3 μM actin (10% ATTO488 labelled) on Lpd^850–1250^(44A)-derivatized, SNAP-Capture magnetic beads in the presence of 5 µM Pfn1 and 50 nM CP after pre-incubation with soluble mVASP variants. The beads were additionally functionalized with trace amounts (5%) of ATTO647-labelled GST-SNAP to enable their visualization. Multi-color TIRF images were captured in 5 sec intervals for a period of 20 min. Time is indicated in min:sec. Scale bar, 10 µm.

**Movie 3: Loss of Ena/VASP impairs *Listeria* actin comet-tail formation.** Representative epifluorescence and phase-contrast, time-lapse movies of *Listeria*-infected B16-F1 cells expressing low levels of EGFP-tagged mVASP-WT together with mScarlet3-actin during migration on laminin. Infected EVM-KO cells expressing mScarlet3-actin alone served as negative control. In agreement with previous studies ^63^, VASP showed a pronounced enrichment at *L. monocytogenes* poles constituting surfaces for active actin assembly and thus the formation of comet tails pushing the bacteria through the cytosol. By contrast, EVM-KO cells failed to support robust comet tail formation, but instead generated only short actin filament stubs at bacterial surfaces mediating significantly reduced, bacterial motility. For bacterial speed determination in these and other conditions, see Supplementary Fig. 5. Time is indicated in min:sec. Scale bar, 10 µm.

**Movie 4: Phosphomimetic mVASP-Y16E impairs *Listeria* actin comet-tail formation in EVM-KO cells.** Representative epifluorescence and phase-contrast, time-lapse movies of *Listeria*-infected B16-F1 cells co-expressing low levels of mScarlet3-actin with either EGFP-mVASP-WT (top) or EGFP-mVASP-Y16E (bottom) during their migration on laminin. Note the clear suppression of prominent actin comet tail formation in cells rescued with phosphomimetic mVASP-Y16E, as opposed to wild-type VASP. Time is indicated in min:sec. Scale bar, 10 µm.

**Movie 5: Phosphomimetic UNC-34-Y17E phenocopies the UNC34-null phenotype.** Representative brightfield time-lapse movies of the wild-type, UNC-34 null, phosphomimetic UNC-34-Y17E and nonphosphorylatable UNC-34-Y17F L4 larvae migrating on bacterial lawns. Note the reduced body size and uncoordinated locomotion of UNC-34-null and UNC-34-Y17E animals. Time is indicated in sec. Scale bar, 5 mm.

## Notes

### Competing Interest Statement

The authors have declared no competing interest.

